# Functional divergence of D-, A-, and B-type cyclins in the moss *Physcomitrium patens*

**DOI:** 10.64898/2026.11.22.768468

**Authors:** Yirong Gan, Jingtong Ruan, Mengying Li, Chengxiang Li, Peishan Yi

## Abstract

Cyclins and cyclin-dependent kinases (CDKs) are central cell-cycle regulators that have undergone diversification in land plants; however, the functional implications of this divergence remain incompletely understood, particularly in non-vascular lineages. Here, we systematically dissected the roles of the three principal cyclin classes—D, A, and B— in the model moss *Physcomitrium patens*. We show that CYCD2 exhibits constitutive expression throughout the cell cycle, CYCA2 is degraded during mitosis and transcriptionally activated post-mitotically, whereas CYCB1 is M-phase-specific and rapidly depleted at anaphase. Overexpression and loss-of-function analyses revealed functional divergence among these classes: cyclin D primarily governs cell cycle entry and vegetative growth; cyclin B regulates mitotic progression with its overexpression causing division failure and arresting gametophore development; cyclin A regulates S-to-G2/M transition and exhibits overlapping functions with both D- and B-type cyclins, contributing to vegetative growth, mitotic fidelity, and gametophore development. Yeast two-hybrid assays showed that CYCB2 associates broadly with both CDKAs and CDKBs, CYCA2 preferentially interacts with CDKAs, and CYCD1 binds CDKAs along with a subset of CDKBs. Collectively, our results reveal both overlapping and class-specific contributions of D-, A-, and B-type cyclins to cell-cycle progression and development in *P. patens*, and suggest that complex tissues (gametophores) are particularly vulnerable to mitotic cyclin dysfunction. These findings provide a foundation for understanding how cyclin diversification underpins developmental plasticity across different contexts in non-vascular land plants.

**Significance statement:** The cyclin-CDK machinery has diverged in land plants, yet the functional significance of this divergence remains incompletely understood. By systematically dissecting D-, A-, and B-type cyclins in the moss *Physcomitrium patens*, this study uncovers both overlapping and class-specific roles in cell-cycle control, and shows that complex tissues are disproportionately vulnerable to mitotic cyclin dysfunction. These findings provide a framework for understanding how cyclin specialization supports distinct developmental contexts in a moss.

## Introduction

In eukaryotes, the cell division cycle is universally driven by cyclin/cyclin-dependent kinase (CDK) complexes (Harashima et al. 2013, Shimotohno et al. 2021). While this core mechanism is broadly conserved, plant cyclin/CDK complexes possess several distinctive features. Notably, plants lack E-type cyclins and have evolved a plant-specific CDK family, CDKB (Vandepoele et al. 2002). In addition, plants have experienced a notable expansion of cyclins and CDKs. For instance, *Arabidopsis* encodes 49 cyclins and 29 CDKs, whereas 29 cyclins and 20 CDKs are present in humans (Renaudin et al. 1996, Burssens et al. 1998, Joubes et al. 2000, Vandepoele et al. 2002, Wang et al. 2004, Cao et al. 2014, Shimotohno et al. 2021). This expansion appears to have occurred late in land plant evolution, as green algae and bryophytes possess fewer cyclins, CDKs, and other core cell-cycle regulators (Bisova et al. 2005, Romani et al. 2026).

In plants, CDKA and CDKB are the two major CDKs that regulate the cell cycle. These two subfamilies differ in their cyclin-binding motifs: CDKA contains the canonical PSTAIRE motif, whereas CDKB carries a modified PPT(A/T)LRE motif (Ferreira et al. 1991, Hirayama et al. 1991, Imajuku et al. 1992, Burssens et al. 1998, Joubes et al. 2000, Shimotohno et al. 2021). CDKBs are unique to the green lineage (Joubes et al. 2000). CDKB homologs in green algae, but not in land plants, complement *cdc2/cdc28* in yeast (Ferreira et al. 1991, Hirayama et al. 1991, Imajuku et al. 1992, Corellou et al. 2005, Cizkova et al. 2008), suggesting that CDKBs underwent functional divergence around land colonization or earlier. CDKAs and CDKBs both govern mitotic progression. However, unlike CDKBs, which play a specific role in mitosis (Porceddu et al. 2001, Andersen et al. 2008, Nowack et al. 2012), CDKAs additionally regulate G1/S transition (Nowack et al. 2012, Zhao et al. 2012).

As in animals, plant A-type and B-type cyclins (CYCAs and CYCBs) primarily regulate the mitotic phase (Hemerly et al. 1992, Menges et al. 2005, Boudolf et al. 2009); D-type cyclins (CYCDs) are crucial for cell cycle entry and G1/S transition (Cockcroft et al. 2000, Boniotti and Gutierrez 2001, Nakagami et al. 2002, Schnittger et al. 2002, Menges et al. 2006). The expression and function of CYCBs are largely restricted to the mitotic stage (Hemerly et al. 1992, Setiady et al. 1995, Fuerst et al. 1996, Reichheld et al. 1996, Ito et al. 1997, Menges et al. 2005), whereas members of the CYCD and CYCA families can have divergent functions. For instance, *Arabidopsis* AtCYCD3;1 and AtCYCD4;1 atypically control DNA synthesis and G2/M progression (Schnittger et al. 2002, Kono et al. 2003); tobacco NtCYCA3;2 is involved in G1/S transition and DNA replication (Yu et al. 2003). Accordingly, members of CYCAs and CYCDs display varied expression patterns (Setiady et al. 1995, Fuerst et al. 1996, Reichheld et al. 1996, Ito et al. 1997, Menges et al. 2005). In *Arabidopsis*, AtCYCA1;1 and AtCYCA2;1-2;3 peak at M phase, AtCYCA3;1 and AtCYCA3;2 peak at S phase, and AtCYCA1;2 remains constant throughout the cell cycle (Menges et al. 2005). S-phase expression of CYCA members has also been observed in tobacco and *Catharanthus roseus* (Setiady et al. 1995, Reichheld et al. 1996). Similarly, although some CYCDs show classical enrichment at G1 (AtCYCD5;1 and AtCYCD3;3), others are found to peak at S (AtCYCD4;1 and AtCYCD4;2) or G2/M (AtCYCD3;1, NtCYCD2;1, and NtCYCD3;1), or remain steady during the cell cycle (AtCYCD3;2 and AtCYCD2;1) (Fuerst et al. 1996, Sorrell et al. 1999, Menges et al. 2005). These divergences are considered critical for cell cycle adaptation and plant plasticity during developmental and environmental responses (Shimotohno et al. 2021).

The expression of cyclins is tightly controlled at both transcriptional and post-translational levels. Particularly, mitotic cyclins are typically expressed at G2 phase and are degraded through the anaphase-promoting complex/cyclosome (APC/C) (Fung and Poon 2005, Berckmans and De Veylder 2009, Shimotohno et al. 2021). APC/C is an E3 ligase that recognizes destruction boxes (D-boxes) in mitotic cyclins and targets them to the 26S proteasome in anaphase (Pines 2011, Willems and De Veylder 2022). In mammals, the transcription of mitotic cyclins depends on B-Myb and Forkhead box M1 (FoxM1) transcription factors scaffolded by the MuvB complex (Fischer et al. 2022). In plants, the MYB3R family fulfils this role through direct binding to M-specific activator (MSA) cis-elements in the promoters of mitotic cyclins and other M-phase-specific targets (Ito et al. 1998, Ito et al. 2001, Haga et al. 2007). This mechanism appears to be widely conserved, as evidenced by functional studies of MYB3Rs and the presence of MSA elements in many G2/M-phase genes across distinct species (Ito et al. 1998, Ito et al. 2001, Haga et al. 2007). Other transcription factors, such as TCP20, also contribute to G2/M-phase expression of CYCBs and can enhance MSA-mediated transcription (Li et al. 2005).

As sister clades of early-diverging land plants, bryophytes serve as valuable models for investigating the functional divergence of cyclins and CDKs. The moss *Physcomitrium patens* (*P. patens*) and the liverwort *Marchantia polymorpha* (*M. polymorpha*) have been established as non-vascular models (Naramoto et al. 2022). *M. polymorpha* retains a minimal set of core cell cycle regulators (Chen et al. 2025). Some components such as CYCAs and CDKBs are expanded in moss (Chen et al. 2025, Romani et al. 2026). Heterologous complementation assays in yeast suggest that *P. patens* CDKAs and CYCD1 play conserved roles in G1/S transition (Lorenz et al. 2003, Bao et al. 2022). CYCD members in moss are also proposed to partner with distinct CDKAs and differentially regulate the G1/S and G2/M transitions, respectively (Peramuna et al. 2023). However, CDKAs are dispensable for gametophyte development, although G1/S transition is delayed in their knockout mutants (Bao et al. 2022). Interestingly, CDKAs are essential for phototropism and chloroplast photo-relocation (Bao et al. 2022). In addition, *P. patens* possesses a remarkable capacity to regenerate following physical or DNA damage (Ishikawa and Hasebe 2022). During regeneration, CYCD1 forms a complex with CDKAs to promote cell cycle re-entry (Ishikawa et al. 2011). Despite these progresses, how cyclins and CDKs differentially coordinate cell cycle progression and development in *P. patens* remains largely unexplored. To address this, we performed comprehensive analyses of three major cyclin classes—D-, A-, and B. Our results indicate that cyclin D and cyclin B predominantly regulate the G1 and M phases, respectively, whereas cyclin A functions throughout the cell cycle. At the tissue level, cyclin D moderately regulates protonema and gametophore growth; cyclin B dysregulation severely impairs gametophore formation; cyclin A plays pleiotropic roles in protonema growth and gametophore emergence and development. Moreover, these cyclin types differentially interact with CDKA and CDKB members. Collectively, we show that D-, A-, and B-type cyclins have functionally diverged in *P. patens* and play important roles in cell-cycle progression and development.

## Results

### The *P. patens* genome encodes a conserved set of cyclin/CDK components

We systematically analyzed all cyclins and CDKs in the moss *P. patens* genome and performed phylogenetic analysis with homologs in the liverwort *M. polymorpha* and flowering plant *Arabidopsis* (Table S1). Consistent with earlier findings (Renaudin et al. 1996, Burssens et al. 1998, Joubes et al. 2000, Vandepoele et al. 2002, Wang et al. 2004, Shimotohno et al. 2021, Chen et al. 2025, Romani et al. 2026), most of these core regulators are conserved in the three species (Figure 1). The SOLO DANCERS (SDS)-type cyclin, which is required for homologous recombination during meiosis in *Arabidopsis* (Azumi et al. 2002), is absent in both bryophytes. Several cyclins were found only in bryophytes including *P. patens*, *M. polymorpha*, and the moss *Ceratodon purpureus*, but were not identified in other plant lineages via BLAST searches in the UniProt and Phytozome databases; we therefore designated these as bryophyte-specific cyclins (cyclin BSs, Figure 1). The *P. patens* genome encodes ten CYCAs and two CYCBs. D-box motifs are present in all moss CYCAs and CYCBs but are absent in AtCYCA3;3, AtCYCB2;5, and MpCYCB2 (Figure S1). Interestingly, moss CYCAs fall into two subclades: one (CYCA4 to CYCA9) contains a well-conserved D-box (RX_2_LX_4_N), whereas the other features an atypical RX_2_LX_4_K motif with Gly and Pro at the -2 and +3 positions, respectively (Figures 1 and S1a). In *M. polymorpha* and *P. patens*, all CYCAs are predicted to contain at least one nuclear localization signal (NLS) upstream of the D-box, whereas this signal is present only in six out of the ten CYCAs in *Arabidopsis* (Figure S1a).

**Figure 1.**
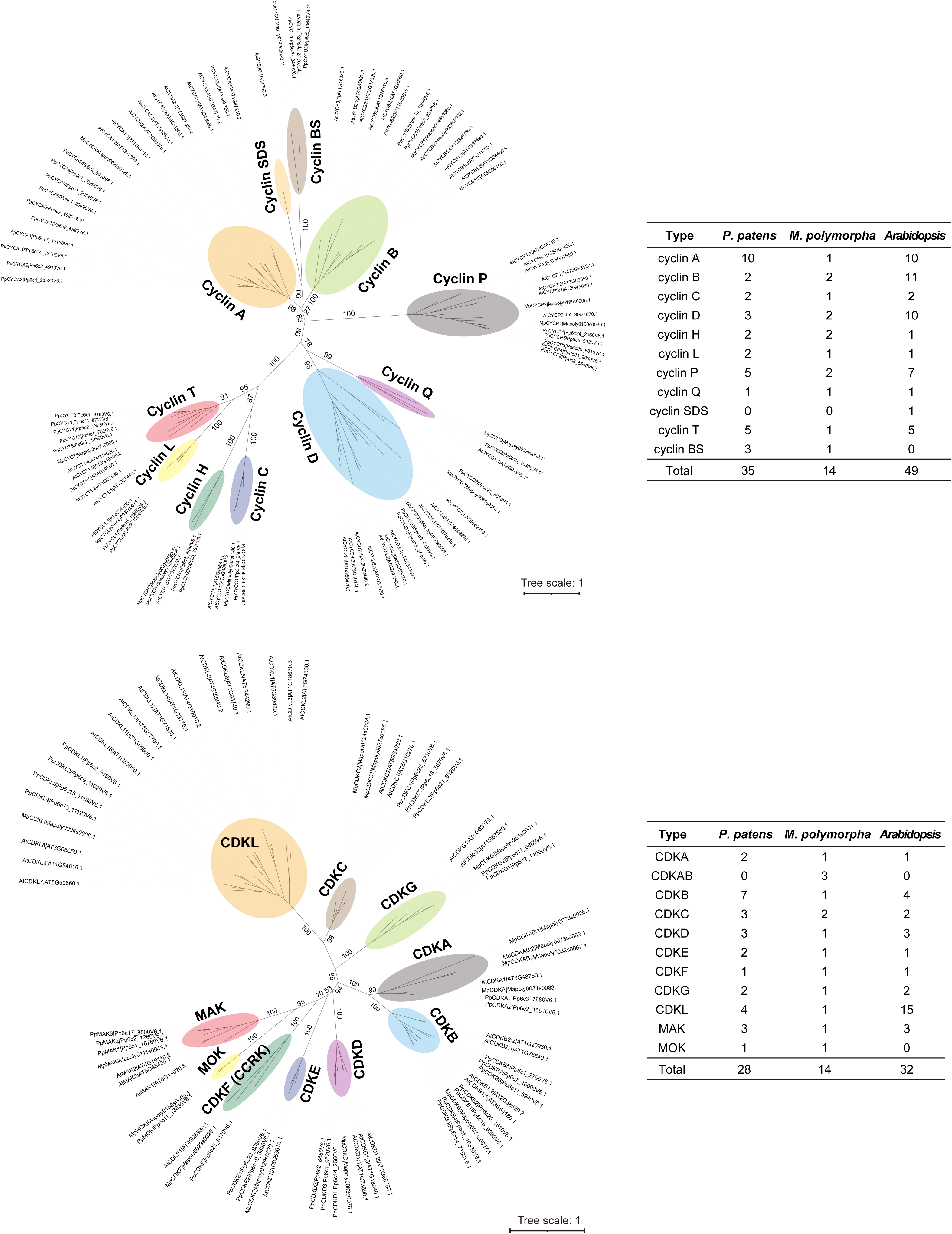
Phylogenetic classification of cyclin and CDK family members in *P. patens*. Classification is based on references (Vandepoele et al. 2002, Wang et al. 2004, Shimotohno et al. 2021, Romani et al. 2026). Cyclin BS, bryophyte-specific cyclin; asterisks indicate proteins with incomplete cyclin domains.

D-type cyclins typically feature an LXCXE motif that mediates binding to Retinoblastoma-related proteins (Dowdy et al. 1993). This motif is conserved in PpCYCD1/2 and MpCYCD1 but is divergent in the phylogenetically distant PpCYCD3 and MpCYCD2 (Figures 1 and S1c). Furthermore, most CYCDs—with the exception of MpCYCD2, PpCYCD3, and the AtCYCD4 group—are predicted to carry a C-terminal NLS immediately downstream of a putative PEST motif (Soni et al. 1995) (Figure S1c). Human cyclin D lacks this NLS but contains a PEST motif region. The corresponding PEST region is relatively conserved in these plant CYCDs. In mammals, PEST motif mediates cyclin D relocation from the nucleus to the cytoplasm and promotes its degradation during S phase (Alt et al. 2000, Simoneschi et al. 2021). These observations imply that plant CYCDs may similarly undergo nucleocytoplasmic shuttling and regulated proteolysis. Indeed, all D-type cyclins in *Arabidopsis* localize to the nucleus, with AtCYCD2;1, AtCYCD4;1, and AtCYCD4;2 also detected in the cytoplasm (Boruc et al. 2010, Strzalka et al. 2015); moreover, several CYCDs, including AtCYCD2;1 and AtCYCD3;1, are subject to proteasome-dependent degradation (Planchais et al. 2004, Sanz et al. 2011).

Similar to *Arabidopsis* (Joubes et al. 2000, Vandepoele et al. 2002, Shimotohno et al. 2021), the CDK family in *P. patens* comprises CDKA through CDKG and a CDK-like (CDKL) group (Figure 1). The CDKA homologs retain the canonical PSTAIRE cyclin-binding motif found in human CDK1 and CDK2 (Meyerson et al. 1992). Among the other groups, CDKB, CDKC, CDKD, CDKE, and CDKG feature an altered P(S/P/A)T(T/A)LRE, PITAIRE, N(F/V/I)TALRE, SPTAIRE, and P(L/M)TSLRE motif, respectively (Figure S1d). Notably, *M. polymorpha* possesses three divergent CDKs distantly related to CDKA (Figure 1). This family, designated CDKAB (Romani et al. 2026), carries a highly modified cyclin-binding motif (Figure S1d). Unlike these CDKs, all CDKFs as well as CDKLs lack a recognizable PSTAIRE motif. The CDKF family is likely a conserved CDK-activating kinase, as has been functionally demonstrated in *Arabidopsis* (Shimotohno et al. 2004). Beyond these core CDKs, we identified two additional groups of cell cycle-related kinases—MALE GERM CELL-ASSOCIATED KINASE (MAK) and MAPK/MAK/MRK OVERLAPPING KINASE (MOK) in *P. patens* and *M. polymorpha* (Figure 1). In green algae and animals, MAK and MOK negatively regulate cilium/flagellum elongation during G0/G1 phase (Berman et al. 2003, Burghoorn et al. 2007, Omori et al. 2010, Broekhuis et al. 2014, Zhang et al. 2025). Our BLAST searches also identified MAK homologs across a broad range of land plants, including the lycophytes *Selaginella moellendorffii* and *Diphasiastrum complanatum*, the fern *Ceratopteris richardii*, the gymnosperm *Thuja plicata*, and the angiosperms *Amborella trichopoda* and *Arabidopsis*. In contrast, MOK homologs were found in lycophytes (*S. moellendorffii* and *D. complanatum*) and ferns (*Ceratopteris richardii* and *Adiantum capillus-veneris*), but were absent from *T. plicata*, *A. trichopoda*, and *Arabidopsis*. Reciprocal BLAST analyses further confirmed MOK’s absence in other gymnosperms, such as *Taxus chinensis* and *Cryptomeria japonica*. These findings suggest that MOK was evolutionarily lost in seed plants and implicate a role of MAK—and potentially MOK in bryophytes and lycophytes—in cellular events during G0/G1 phase. Interestingly, CDKFs are homologous to human CELL CYCLE RELATED KINASE (CCRK)/CDK20 and are conserved in green algae and worms. These kinases regulate cilium/flagellum homeostasis by modulating MAK activity (Tam et al. 2007, Wang and Kung 2012, Yang et al. 2013, Yi et al. 2018, Zhang et al. 2025). Whether CDKFs act together with MAK to control cell-cycle-related processes or exert additional functions remains an open question.

## D-, A-, and B-type cyclins exhibit distinct expression profiles during the cell cycle

We next focused on *CYCD2*, *CYCA2*, and *CYCB1* and analyzed their expression patterns because D-, A-, and B-type cyclins are core regulators of the cell cycle and these genes are highly expressed in actively proliferating filamentous protonemata (Figure S2). To simultaneously visualize transcriptional and translational expression, we employed a self-cleaving F2A peptide strategy (Khosla et al. 2020), which enables translation of two fluorescent protein fusions from a single open reading frame. We first validated F2A functionality by fusing an N-terminal membrane-targeting myristoylation signal to mCherry (Lai et al. 2023), followed by F2A, an SV40 NLS, and Citrine (Figure S3a). Transient expression of this reporter in protoplasts confirmed membrane localization of mCherry and nuclear localization of SV40-Citrine (Figure S3b). Identical patterns were detected with stably transformed lines (Figure S3c). Quantitatively, the majority of Citrine signal (>80%) was concentrated in the nucleus, while no nuclear localization of mCherry was observed, demonstrating the high cleavage efficiency of F2A in moss (Figure S3d).

Using the F2A system, *CYCD2*, *CYCA2*, and *CYCB1* were expressed under the control of their endogenous promoters, with each construct harboring a C-terminal mCherry tag followed by F2A-SV40-Citrine. In this configuration, Citrine fluorescence reports transcriptional expression while mCherry fluorescence indicates translational expression (Figure 2a). As shown in Figures 2b and 2c, all three genes were transcriptionally active in proliferating cells, including protonema tip cells, branch initial cells, gametophore buds, and phyllid initials. At the protein level, both CYCD2-mCherry and CYCA2-mCherry exhibited nuclear localization in pre-mitotic and post-mitotic cells (Figures 2d and 2e, Movies S1 and S2), similar to their *Arabidopsis* homologs (Boruc et al. 2010, Strzalka et al. 2015). Notably, CYCA2 fluorescence decreased by approximately 40% in post-mitotic cells, whereas the CYCD2 signal remained unchanged (Figure 2f). These findings suggest that CYCA2, unlike CYCD2, undergoes regulated degradation during M phase. A truncated variant comprising the N-terminal D-box and NLS of CYCA2 (CYCA2(NT), residues 1-203) recapitulated the expression pattern of the full-length protein (Figures 2g, S4a and S4b, Movie S3). During telophase, the Citrine signal for *CYCD2* progressively increased within the reforming nucleus, peaking approximately 70 minutes after anaphase, whereas no change was detected for CYCD2-mCherry (Figure 2h). The Citrine signal for CYCA2(NT) continued to rise until 120 minutes, with CYCA2(NT)-mCherry fluorescence concurrently increasing and ultimately reaching pre-mitotic levels (Figures 2h and S4c). Given the high stability of the Citrine fluorophore, the early post-anaphase increase (within 70 minutes) in nuclear signal likely reflects nuclear import of pre-existing SV40-Citrine proteins rather than de novo transcription. Accordingly, we infer that *CYCD2* expression remains largely constant throughout the post-mitotic phase, whereas *CYCA2* transcription is activated post-mitotically.

**Figure 2.**
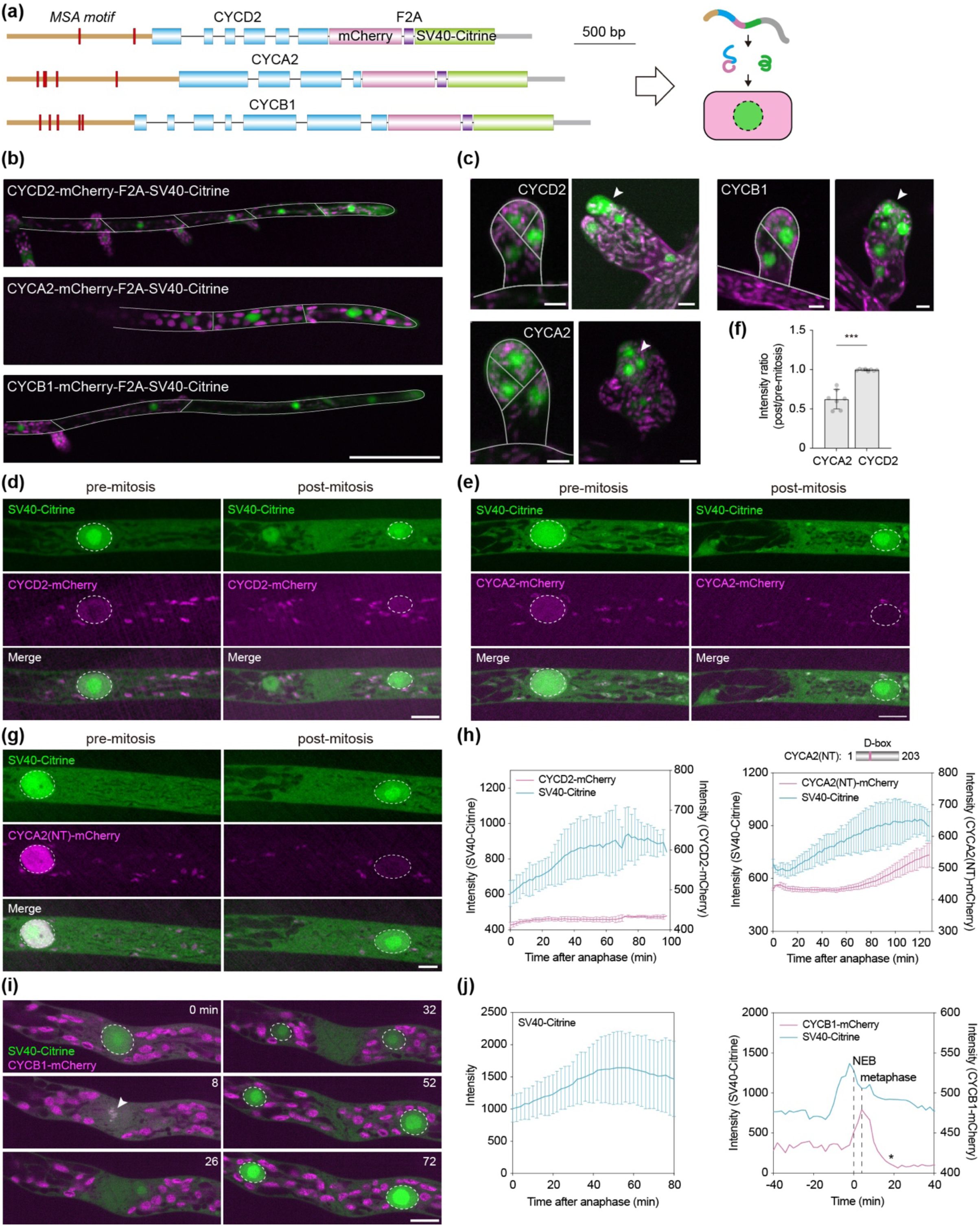
**Expression profiles and subcellular localization of CYCD2, CYCA2, and CYCB1.**(a) Schematic of F2A-mediated dual-reporting constructs for *CYCD2*, *CYCA2*, and *CYCB1*. Promoters and exons are indicated by brown lines and blue boxes, respectively. MSA, M-specific activator cis-element; F2A, self-cleaving peptide; SV40, nuclear localization signal. (b) Expression of *CYCD2*, *CYCA2*, and *CYCB1* in protonema cells. The mCherry signal was undetectable at this magnification. The magenta channel corresponds to chlorophyll autofluorescence. Scale bar, 100 μm. (c) Expression of *CYCD2*, *CYCA2*, and *CYCB1* in gametophore buds. Arrowheads indicate phyllid initials. Scale bars, 10 μm. (d, e) Expression of *CYCD2* (d) and *CYCA2* (e) in pre-mitotic and post-mitotic cells. Nuclei are outlined by dashed circles. Scale bars, 10 μm. (f) Quantification of nuclear fluorescence intensity ratios in post-versus pre-mitotic cells. Data are shown as mean ± SD. ***, *p* < 0.001; Mann-Whitney test. *CYCA2*: n = 7; *CYCD2*: n = 8. (g) Expression of the *CYCA2* N-terminal region (*CYCA2(NT)*) in pre-mitotic and post-mitotic cells. Scale bar, 10 μm. (h) Post-mitotic expression profile of *CYCD2* and *CYCA2(NT)*. Data are shown as mean ± SD. *CYCD2*: n = 4; *CYCA2(NT)*: n = 3. (i) Mitotic expression and localization of *CYCB1*. The arrowhead indicates metaphase chromosomes. Scale bar, 10 μm. (j) Expression profile of *CYCB1* at post-mitotic (left) and mitotic (right) stages. Note the sharp decline in CYCB1 signal upon anaphase onset (asterisk). Data in the left panel are shown as mean ± SD (n = 4).

In contrast to CYCD2 and CYCA2, CYCB1-mCherry was not visible in the nucleus; instead, it was recruited to the nuclear envelope in prophase and associated with chromosomes upon nuclear envelope breakdown (Figures 2i and 2j, Movie S4). During anaphase, CYCB1-mCherry fluorescence was depleted within 20 minutes, suggesting its rapid degradation. The Citrine signal for *CYCB1* followed a profile analogous to that of *CYCD2* (Figure 2j). We assessed whether the N-terminal domain of CYCB1 (CYCB1(NT), residues 1-251) is sufficient to mediate mitotic turnover. VPS4 is a key endosome biogenesis factor (Ruan et al. 2026). It localizes to late endosomes and the cell plate during cytokinesis (Figures S5a and S5b, Movie S5). When fused with CYCB1(NT), VPS4 was redirected to chromosomes, mirroring CYCB1 localization. During anaphase, its signals on endosomes and chromosomes were both rapidly depleted (Figure S5c, Movie S6). These data show that CYCB1(NT) is sufficient to confer cell-cycle-dependent localization and dynamics.

Altogether, our findings suggest that *CYCD2* expression remains constant throughout the cell cycle, *CYCA2* is transcriptionally activated early in the cell cycle, and *CYCB1* is a M-phase-specific cyclin. Furthermore, both CYCA2 and CYCB1 are downregulated in mitosis presumably through D-box-mediated protein degradation.

## D-, A-, and B-type cyclins negatively regulate protonema cell growth

When cultured on standard BCDAT medium, the *CYCD2*, *CYCA2*, and *CYCB1* overexpression lines displayed varied developmental phenotypes. Notably, colony growth was markedly inhibited by *CYCA2* and *CYCD2* overexpression, whereas no significant changes were observed in *CYCB1* overexpressors (Figures 3a and 3b). Accordingly, protonema cells were substantially shortened in *CYCD2* and *CYCA2* overexpressors, but remained unaffected by *CYCB1* overexpression (Figures 3c and 3d). Cell width was diminished and increased in *CYCA2* and *CYCB1* overexpressors, respectively (Figures 3c and 3d). These findings indicate that CYCA2 and CYCD2 negatively regulate cell growth, whereas CYCB1 plays a limited role in protonema development.

**Figure 3.**
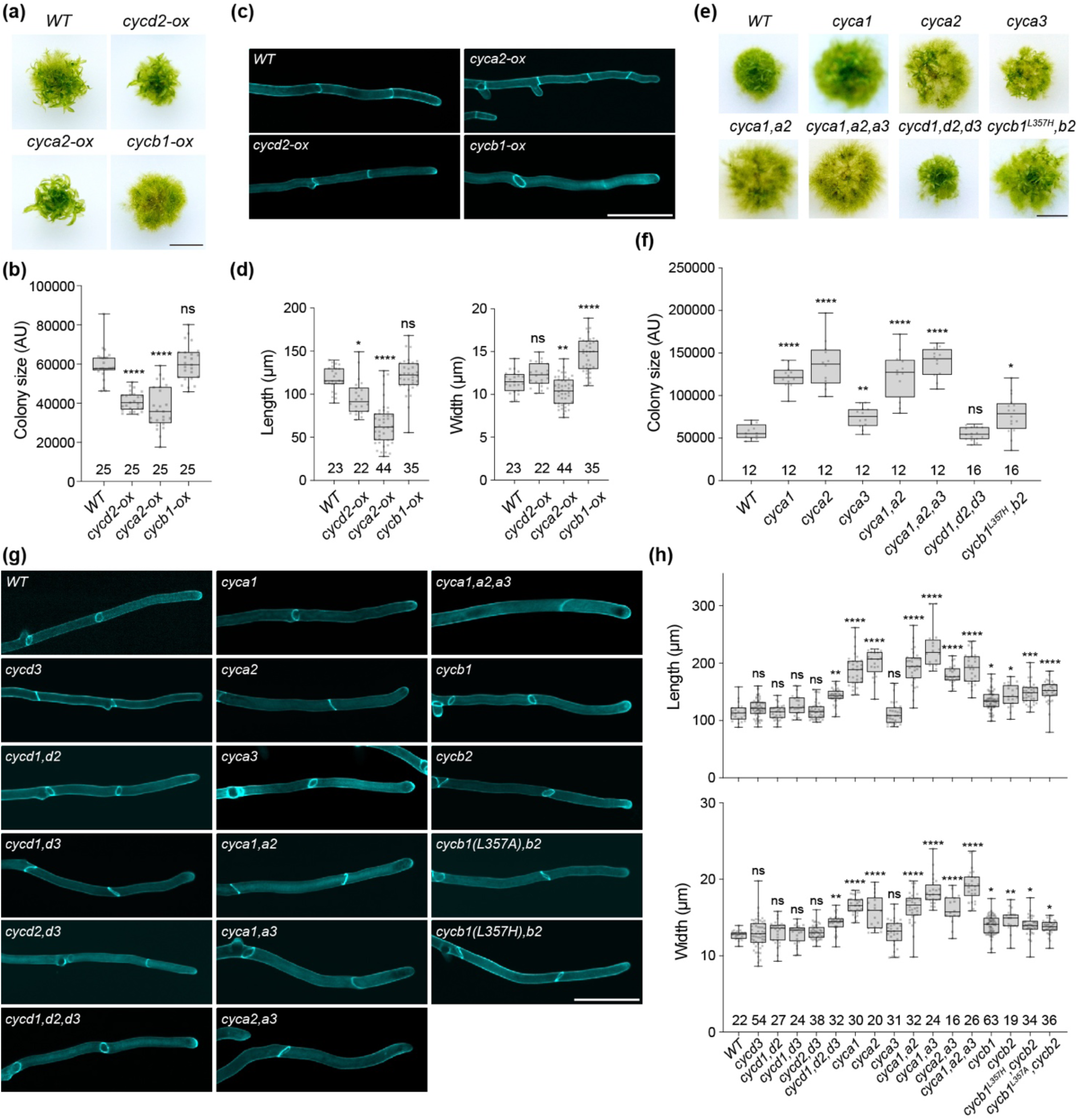
D-, A-, and B-type cyclins negatively regulate protonema growth. (a) Colony morphology of wild-type (*WT*) and *CYCD2-*, *CYCA2*-, and *CYCB1*-overexpression lines. Scale bar, 5 mm. (b) Quantification of colony sizes in overexpression lines. Data are shown with box plots, displaying the interquartile range, the median, and minimum and maximum values. Sample sizes are indicated below the bars. Statistical significance relative to *WT* was assessed using Welch’s ANOVA followed by Dunnett’s T3 post hoc multiple-comparison test; ns, not significant; ****, *p*<0.0001. (c) Protonema cell morphology in the overexpression lines, stained with calcofluor white. Scale bar, 100 μm. (d) Quantification of cell length and width in overexpression lines. Sample sizes are indicated below the bars. Statistical significance relative to *WT* was assessed using Kruskal–Wallis followed by Dunn’s post hoc multiple-comparison test and Welch’s ANOVA followed by Dunnett’s T3 post hoc multiple-comparison test for analyzing cell length and width, respectively; ns, not significant; *, *p*<0.05; **, *p*<0.01; ****, *p*<0.0001. (e) Colony morphology of *cycd*, *cyca*, and *cycb* mutants. Scale bar, 5 mm. (f) Quantification of mutant colony sizes. Sample sizes are indicated below the bars. Statistical significance relative to *WT* was assessed using Welch’s ANOVA followed by Dunnett’s T3 post hoc multiple-comparison test; ns, not significant; *, *p*<0.05; **, *p*<0.01; ****, *p*<0.0001. (g) Protonema cell morphology in mutant lines. Scale bar, 100 μm. (h) Quantification of cell length and width in mutant plants. Sample sizes are indicated below the bars. Statistical significance relative to *WT* was assessed using Kruskal–Wallis followed by Dunn’s post hoc multiple-comparison test; ns, not significant; *, *p*<0.05; **, *p*<0.01; ***, *p*<0.001; ****, *p*<0.0001.

To further interrogate cyclin function, we generated knockout mutants for *CYCD*, *CYCA*, and *CYCB* using CRISPR/Cas9-assisted genome editing (Collonnier et al. 2017). Given that *CYCA1*, *CYCA2*, and *CYCA3* exhibited the highest expression levels among CYCAs in protonemata (Figure S2a), we targeted these three CYCAs, and all CYCDs and CYCBs. We obtained frameshift single mutants as well as various high-order mutant combinations (Figures S6a and S6b). As shown in Figures 3e and 3f, colony growth was significantly enhanced in the *cyca1* or *cyca2* single mutants, but remained unaltered in the *cyca3 single mutant*. This phenotype was further accentuated in the *cyca1,a2* double and *cyca1,a2,a3* triple mutants. Correspondingly, protonema cells in these mutants exhibited increases in both cell length and width (Figures 3g and 3h). In the case of CYCDs, the *cycd3* single and *cycd* double mutants displayed no discernible growth defects; however, cell length and width were increased in the *cycd1,d2,d3* triple mutant (Figures 3g and 3h). These data corroborate the inhibitory roles of CYCDs and CYCAs in protonema cell growth.

Although we successfully isolated single knockout mutants for *cycb1* and *cycb2*, we failed to recover a double knockout line. Nevertheless, both single mutants exhibited slight but statistically significant increases in protonema cell length and width (Figures 3g and 3h). To further dissect CYCB function, we introduced point mutations at Leu357 of *cycb1* in the *cycb2* knockout background by co-transforming degenerate oligonucleotide templates along with CRISPR/Cas9 plasmids (Yi and Goshima 2020) (Figures S6a-S6c). This strategy was anticipated to randomly introduce L357-to-A/D/G or L357-to-P/H/R substitutions, potentially disrupting the hydrophobic interactions between α6 and α7/α8 (Figure S6d). After several trials of transformation, we only obtained L357A and L357H variants. These observations suggest that CYCBs are essential for viability and that substitution of Leu357 with structurally divergent residues, such as Asp, Gly, Pro, and Arg, is likely lethal. Nevertheless, both viable mutants displayed increased cell length, yet cell width remained less affected (Figures 3g and 3h). To determine whether the growth defects arise from altered cell expansion, we quantified growth rates of protonema tip cells in cyclin overexpressors and high-order mutants. However, no significant difference was detected for any of the examined lines (Figure S7). These results suggest that cyclins inhibit protonema cell growth likely through cell cycle regulation rather than by influencing intrinsic cell-expansion capacity.

## Perturbed cyclin expression and function disrupts gametophore development

We next examined cyclin function during gametophore development, a complex three-dimensional tissue that derives from a single side-branch cell on the filamentous protonemata (Moody 2022). Because cytokinin stimulates gametophore initiation (Brandes and Kende 1968), we tested whether cytokinin induces cyclin expression. Consistent with recent single-nucleus RNA-sequencing results (Hata et al. 2025), expression of all selected D-, A-, and B-type cyclin members was upregulated by 2-isopentenyladenine (2iP), with *CYCB1* and *CYCD3* showing the strongest induction (Figure 4a), implicating these cyclins in gametophore development.

**Figure 4.**
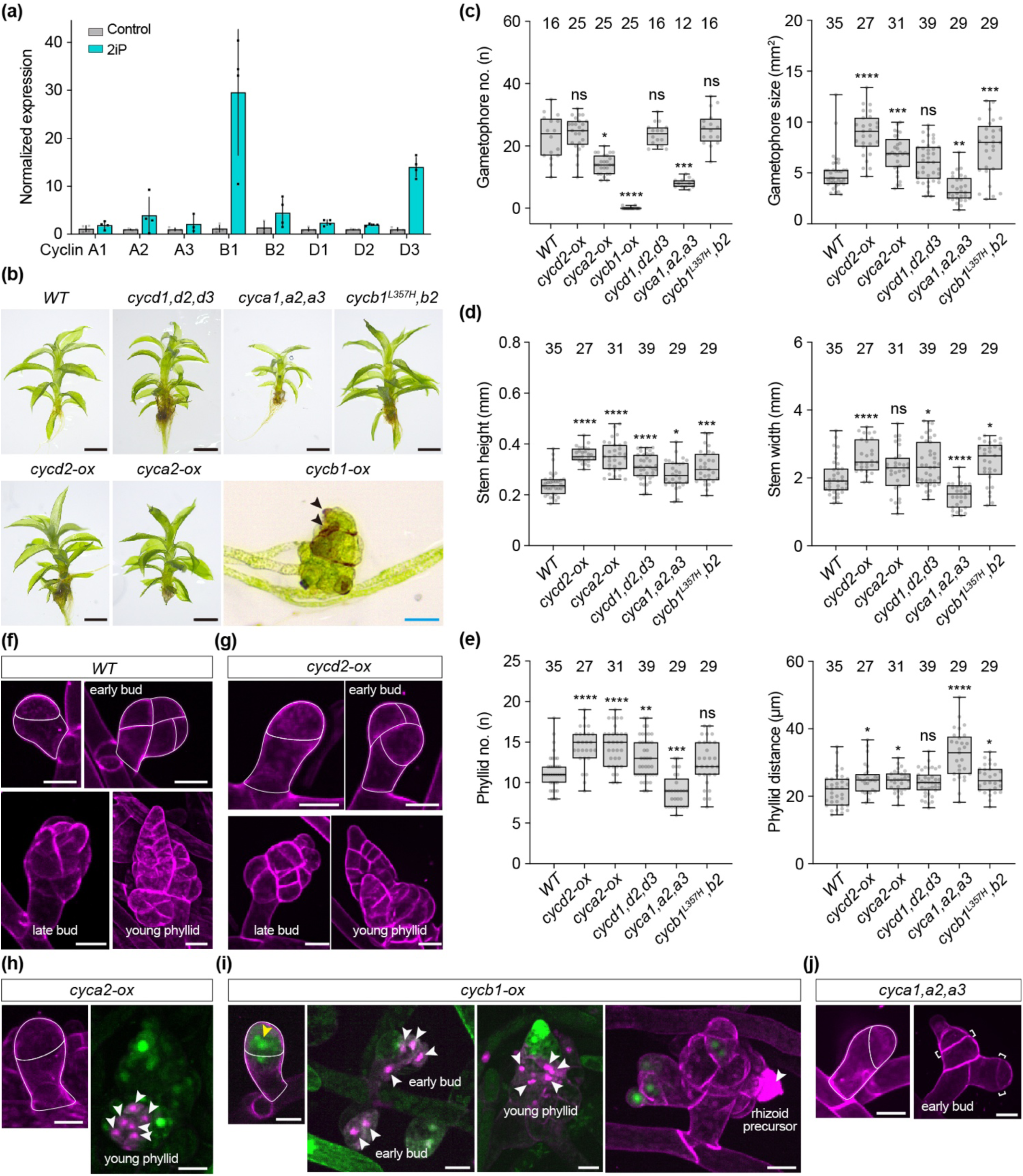
Overexpression and loss-of-function of D-, A-, and B-type cyclins disrupts gametophore development. (a) Transcript levels of the indicated cyclins following treatment with the cytokinin 2iP. Data were normalized to endogenous *EF1α* and are presented as mean ± SD (n = 4). (b) Gametophore morphology of *WT*, cyclin mutants, and cyclin overexpression lines. Scale bars: 1 mm (black) and 0.1 mm (blue). (c-e) Quantification of gametophore number and overall size (c), stem height and width (d), and phyllid number and internodal spacing (e) for the indicated genotypes. Data are shown with box plots, displaying the interquartile range, the median, and minimum and maximum values. Sample sizes are shown above the bars. Statistical significance relative to *WT* was assessed using Kruskal–Wallis followed by Dunn’s post hoc multiple-comparison test for c (left); Welch’s ANOVA followed by Dunnett’s T3 post hoc multiple-comparison test for c (right), d, and e (right); and one-way ANOVA followed by Dunnett’s post hoc multiple-comparison test for e (left). ns, not significant; *, *p*<0.05; **, *p*<0.01; ***, *p*<0.001; ****, *p*<0.0001. (f-j) Propidium iodide (PI) staining (red) of gametophore buds and young gametophores in *WT* (f), *cycd2-ox* (g), *cyca2-ox* (h), *cycb1-ox* (i), and *cyca1,a2,a3* (j) lines. Green fluorescence in (h) and (i) shows SV40-Citrine signal expressed in the corresponding overexpression lines. White arrowheads indicate PI-positive dead cells; yellow arrowhead denotes a binucleate cell in the *cycb1-ox* line; brackets delineate aberrant division planes in the *cyca1,a2,a3* mutant. Scale bars: 20 μm.

Given the functional redundancy among cyclin members (Figure 3), we assessed gametophore phenotypes in the *cycd1,d2,d3* and *cyca1,a2,a3* triple mutants, the *cycb1^L357H^,b2* double mutant, and the *cycd2-ox*, *cyca2-ox*, and *cycb1-ox* overexpression lines. While *cycd2-ox* and knockout of D-type cyclins did not impair gametophore formation, gametophore size increased significantly in *cycd2-ox* and slightly in the knockout lines (Figures 4b and 4c), suggesting that proper D-type cyclin dosage is required for normal morphogenesis. Accordingly, both *cycd2-ox* and *cycd1,d2,d3* plants exhibited increased stem height and width, along with greater phyllid number and spacing, although with the mutant line displaying a milder effect (Figures 4d and 4e). In contrast, gametophore emergence was markedly reduced in both *cyca2-ox* and *cyca1,a2,a3* lines. Furthermore, gametophore size increased in the overexpressor and decreased in the mutant (Figures 4b and 4c). These results indicate that cyclin A plays dual roles in gametophore emergence and growth. Notably, *cyca1,a2,a3* plants developed longer and narrower stems with fewer phyllids, resulting in significantly increased phyllid spacing (Figures 4d and 4e). The *cyca2-ox* line produced more phyllids and showed enhanced stem elongation (Figures 4d and 4e). Thus, cyclin A appears to stimulate phyllid initiation while requiring strict dosage homoeostasis to balance stem growth as does cyclin D. For B-type cyclins, the *cycb1^L357H^,b2* mutant showed no significant effect on gametophore emergence but exhibited generally enhanced growth (Figures 4c-4e). More strikingly, *cycb1-ox* plants showed near-complete arrest at the late bud stage, with phyllids failing to mature and displaying tissue necrosis (Figures 4b and 4c).

To further characterize the cellular defects underlying these phenotypes, we performed propidium iodide staining on the *cycd2-ox*, *cyca2-ox, cycb1-ox*, and *cyca1,a2,a3* lines. Consistent with the macroscopic observations, *cycd2-ox* gametophores developed normally (Figures 4f and 4g). Overexpression of *cyca2* led to occasional cell death in young phyllids, whereas cell death was frequently observed in *cycb1-ox* plants from early buds through the young phyllid stage (Figures 4h and 4i), corroborating the moderate and severe defects in gametophore development observed for these lines, respectively. Although the *cyca1,a2,a3* mutant did not block bud emergence, it caused aberrant division planes in early buds, which appeared to trigger subsequent growth arrest and led to gametophore reduction (Figure 4j).

Together, our data indicate that cyclin D is dispensable for gametophore formation, but is required for normal morphogenesis. Cyclin A exerts pleiotropic roles in gametophore formation, phyllid development, and stem elongation. Cyclin B modulates gametophore growth and must be strictly regulated during gametophore development. The dosage dependence of cyclins A and B highlights the importance of their precise regulation during complex tissue formation.

## D-, A-, and B-type cyclins exert overlapping yet divergent functions in cell cycle regulation

To dissect the specific roles of D-, A-, and B-type cyclins in cell cycle progression, we analyzed the cell division process using time-lapse imaging. CYCD2 and CYCA2 overexpression, driven by their native promoters, did not strongly perturb mitotic progression. In contrast, approximately 43% of the protonema cells (9/21) overexpressing *CYCB1* exhibited delayed and uncoordinated chromosome segregation (Figure 5a, Movies S7 and S8). This defect led to nuclear membrane reassembly around multiple chromatin clusters, culminating in cytokinesis failure and multinucleation. Even in successfully divided cells, the apical daughter cell occasionally underwent abrupt cell death (Figure 5b). These findings underscore a critical role of cyclin B regulation in mitotic fidelity and cell survival. Unlike the severe mitotic defects observed in *CYCB1* overexpressors, cell division proceeded to completion in *cycd1,d2,d3* triple mutants, *cyca1,a2,a3* triple mutants, and *cycb1^L357H^,b2* double mutants. However, both mitosis and cytokinesis were markedly prolonged in these lines (Figures 5c-5f, Movies S9-S12), indicating that all three cyclin classes are collectively required for timely M-phase progression.

**Figure 5.**
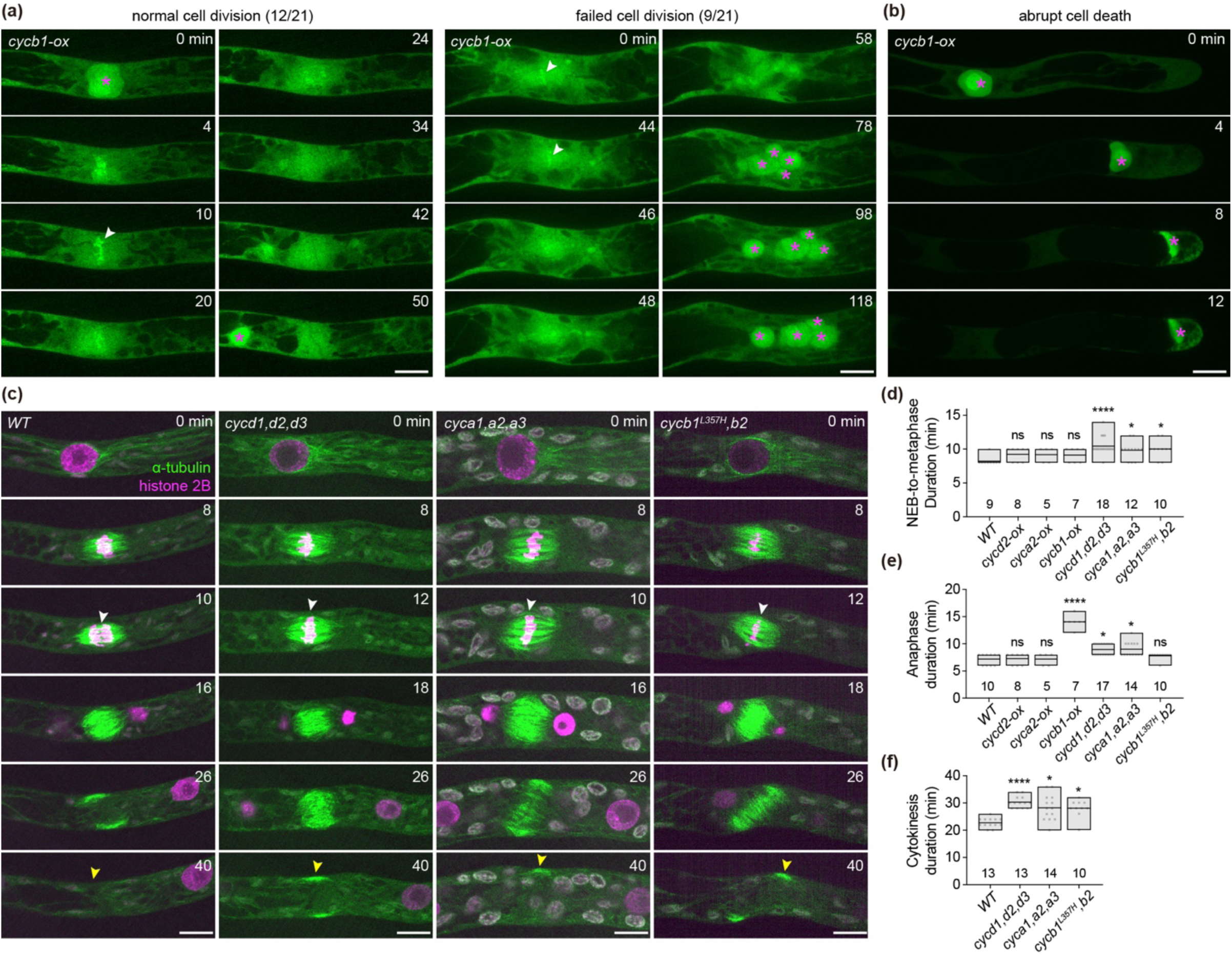
D-, A-, and B-type cyclins are collectively required for cell division progression. (a, b) Representative protonema cells from *CYCB1* overexpressors undergoing normal division (a), displaying mitotic and cytokinesis defects (a), or undergoing cell death (b). Asterisks indicate nuclei; arrowheads mark metaphase chromosomes. Scale bars, 10 μm. (c) Mitotic progression in *WT* and cyclin mutants, visualized with GFP-α-tubulin (green) and histone 2B-RFP (magenta). White arrowheads indicate anaphase onset; yellow arrowheads mark the edges of the phragmoplast. Scale bars, 10 μm. (d-f) Quantification of cell-cycle phase durations. Data are shown with box plots, displaying the mean, and minimum and maximum values. Sample sizes are shown below the bars. Statistical significance relative to *WT* was assessed using Kruskal–Wallis followed by Dunn’s post hoc multiple-comparison test for d and e, and Welch’s ANOVA followed by Dunnett’s T3 post hoc multiple-comparison test for f; ns, not significant; *, *p*<0.05; ****, *p*<0.0001.

To further validate cyclin function, we performed flow cytometry analysis. Although moss gametophytes are haploid, isolated nuclei from both protonemata and gametophores displayed a predominant G2/M (2C) peak, with only a small G1 (1C) population detectable (Figure 6a). A minor 4C peak, likely corresponding to endoreplication cells, was additionally observed in gametophores (Figure 6a). These patterns are consistent with earlier reports that most moss cells are arrested in G2 phase (Schween et al. 2008, Ishikawa et al. 2011, Bao et al. 2022). To further confirm that the major peak indeed represents the 2C population, we used nuclei from *Arabidopsis* flowers as a reference. Consistent with previous findings (Li et al. 2020), *Arabidopsis* nuclei exhibited a major 2C peak alongside endoreplication peaks at 4C and 8C. When nuclei from both species were mixed and co-analyzed, the moss 2C population (∼480 Mb ×2) aligned well with the *Arabidopsis* 8C peak (∼120 Mb ×8) (Figure 6a). The exceptionally small 1C population in moss suggests an abbreviated G1 phase. Consistent with this, endogenously mNG tagged PCNA1 (proliferating cell nuclear antigen 1), a core DNA replication factor, translocated to the nucleus during cytokinesis and subsequently localized to discrete puncta, with fluorescence signal dissipating within 90 minutes (Figure 6b, Movie S13). In *Arabidopsis*, the formation of transient PCNA1 puncta strongly correlates with DNA replication (Yokoyama et al. 2016). Collectively, these data inidcate that moss gametophytes have an exceptionally short G1 phase.

**Figure 6.**
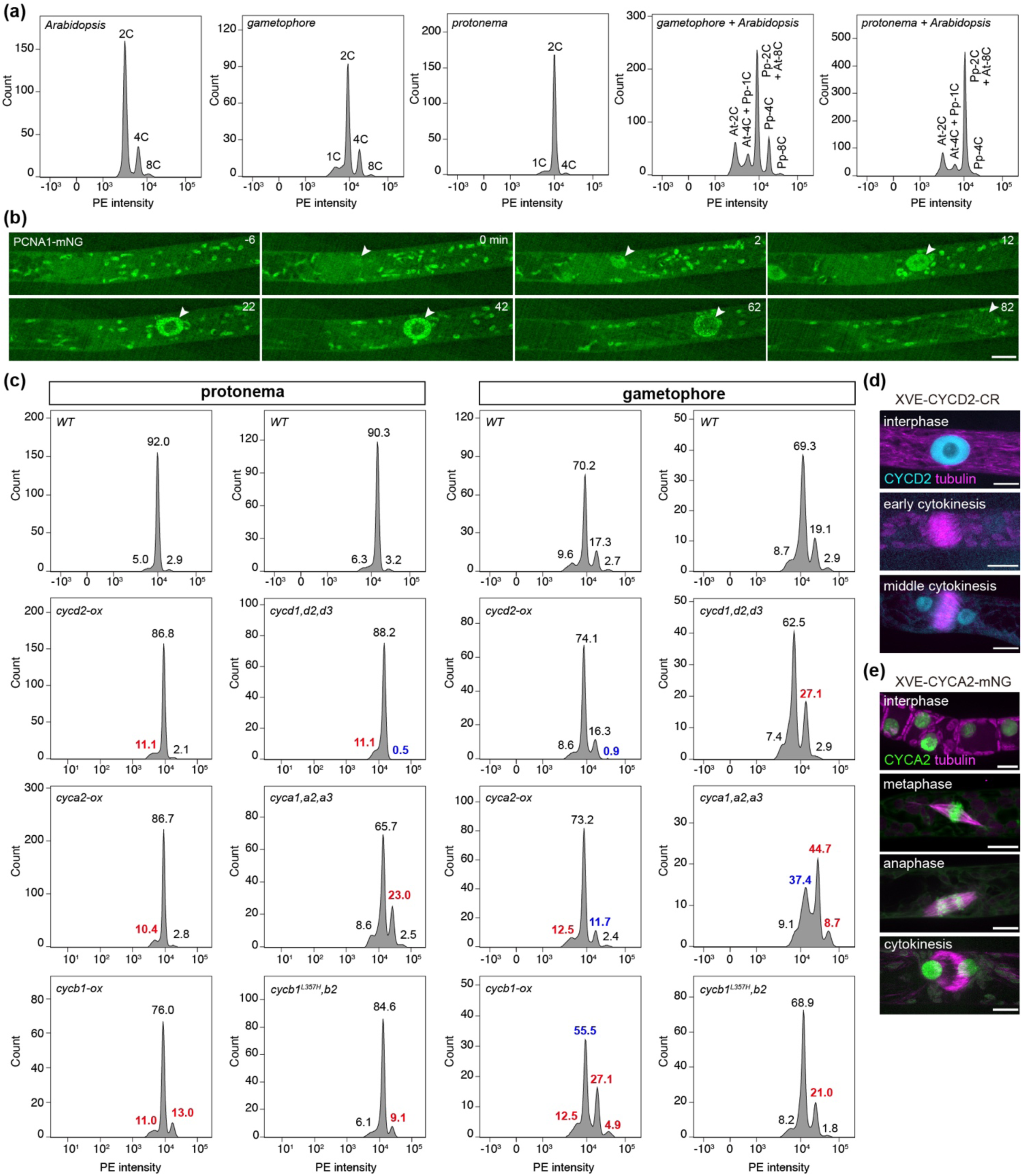
D-, A-, and B-type cyclins are differentially involved in cell cycle regulation. (a) Flow cytometry analysis showed that moss cells are predominantly arrested in G2 phase. (b) Transient nuclear localization of PCNA1 fused with mNeonGreen (mNG) (arrowheads) from the end of anaphase (0 min) to the post-cytokinetic stage. Scale bar, 10 μm. (c) Flow cytometry profiles of cyclin overexpressors and mutants. Populations showing increased or decreased abundance are highlighted in red and blue, respectively. (d, e) Inducible expression of *CYCD2* (d) and *CYCA2* (e) in protonema cells. Microtubules are labeled with mCherry-α-tubulin (magenta). CYCA2 localizes to chromosomes during metaphase and anaphase. XVE, inducible expression cassette comprising the LexA repressor and chimeric VP16-estrogen receptor; CR, Cerulean. Scale bars, 10 μm.

We next examined the cell-cycle profiles of cyclin overexpressors and mutants. In protonema cells, overexpression of *CYCD2*, *CYCA2*, or *CYCB1* all increased the G1 population (Figure 6c), implying that elevated cyclin levels inhibit cell cycle entry. A similar effect was observed for *CYCA2* and *CYCB1* in gametophore cells. Notably, *CYCB1* overexpression also elevated the 2C and 4C populations in protonemata and gametophores, likely due to cytokinesis failure noted earlier (Figure 5b). In *cycd1,d2,d3* triple mutants, the protonemal 2C population decreased, while the 1C population increased, indicating that CYCDs are required for cell cycle entry in this tissue (Figure 6c). Interestingly, in gametophores, loss of CYCDs did not affect cell cycle entry, but instead led to an increase in 4C cells. Conversely, *CYCD2* overexpression reduced the 4C/8C population (Figure 6c). These results suggest that CYCDs either promote M-phase progression or inhibit endoreplication in gametophores. Notably, a marked increase in the 4C population was also observed in both protonema cells and gametophores of the *cyca1,a2,a3* and *cycb1^L357H^,b2* mutants (Figure 6c), implying overlapping functions among these cyclins, although CYCAs and CYCBs appear to have less tissue specificity. Together, these results suggest that CYCDs regulate cell cycle entry in protonemata and all three cyclins may play a role in M-phase progression or endoreplication.

Given that overexpression and mutant lines of cyclin D and cyclin A show similar growth phenotypes (Figures 2d-2h and 3), we asked whether *CYCD2* and *CYCA2* could induce cell division upon ectopic expression. To this end, we employed a β-estradiol-inducible system to drive ubiquitous *CYCD2* or *CYCA2* expression in protonema cells (Kubo et al. 2013). Upon β-estradiol treatment, both proteins were expressed and localized in the nucleus (Figures 6d and 6e). Strikingly, *CYCA2* expression induced mitotic divisions in subapical cells lacking a characteristic pre-mitotic bulge (Figure 6e, observed in 8/26 filaments). Consequently, some subapical cells became severely shortened. In dividing cells, the spindles were unusually narrow, consistent with insufficient microtubule polymerization. Unlike *CYCA2*, we did not observe ectopic subapical cell divisions (28 filaments) upon *CYCD2* expression. Furthermore, ectopically expressed *CYCA2* accumulated on chromosomes (Figure 6e). Such localization was not obvious for endogenously expressed CYCA2 (Movie S2), possibly due to its relatively low expression level and rapid degradation as found for most *Arabidopsis* CYCAs (Boruc et al. 2010). The finding that CYCA2, unlike CYCD2, induces ectopic division implies distinct cell-cycle roles: CYCDs likely act at G1, whereas CYCAs play a role in the S-to-G2/M transition in protonema cells. This interpretation agrees with our flow cytometry data and with the observed post-mitotic upregulation of CYCA2 and PCNA1 (Figures 6b and 6c).

## D-, A-, and B-type cyclins differentially interact with CDKA/B members

Cyclin pairing is critical for CDK function and substrate specificity. To investigate cyclin-CDK associations in moss, we performed yeast two-hybrid assays focusing on D-, A-, and B-type cyclins and CDKA and CDKB members. Notably, transformants expressing CDKA1, CYCA1, CYCA3, and CYCB1 alone yielded few colonies (Figure S8), suggesting that these proteins impede yeast viability possibly due to cell cycle perturbation. Although expression of other cyclin and CDK members alone did not markedly affect yeast viability, approximately 70% of the cyclin-CDKA/B co-transformants (27/39) produced few or no colonies, indicating that these pairs exert additive deleterious effects (Figure 7a). Among the surviving co-transformants, positive interactions were detected for CYCD1-CDKA2 and CYCD2-CDKB6.

**Figure 7.**
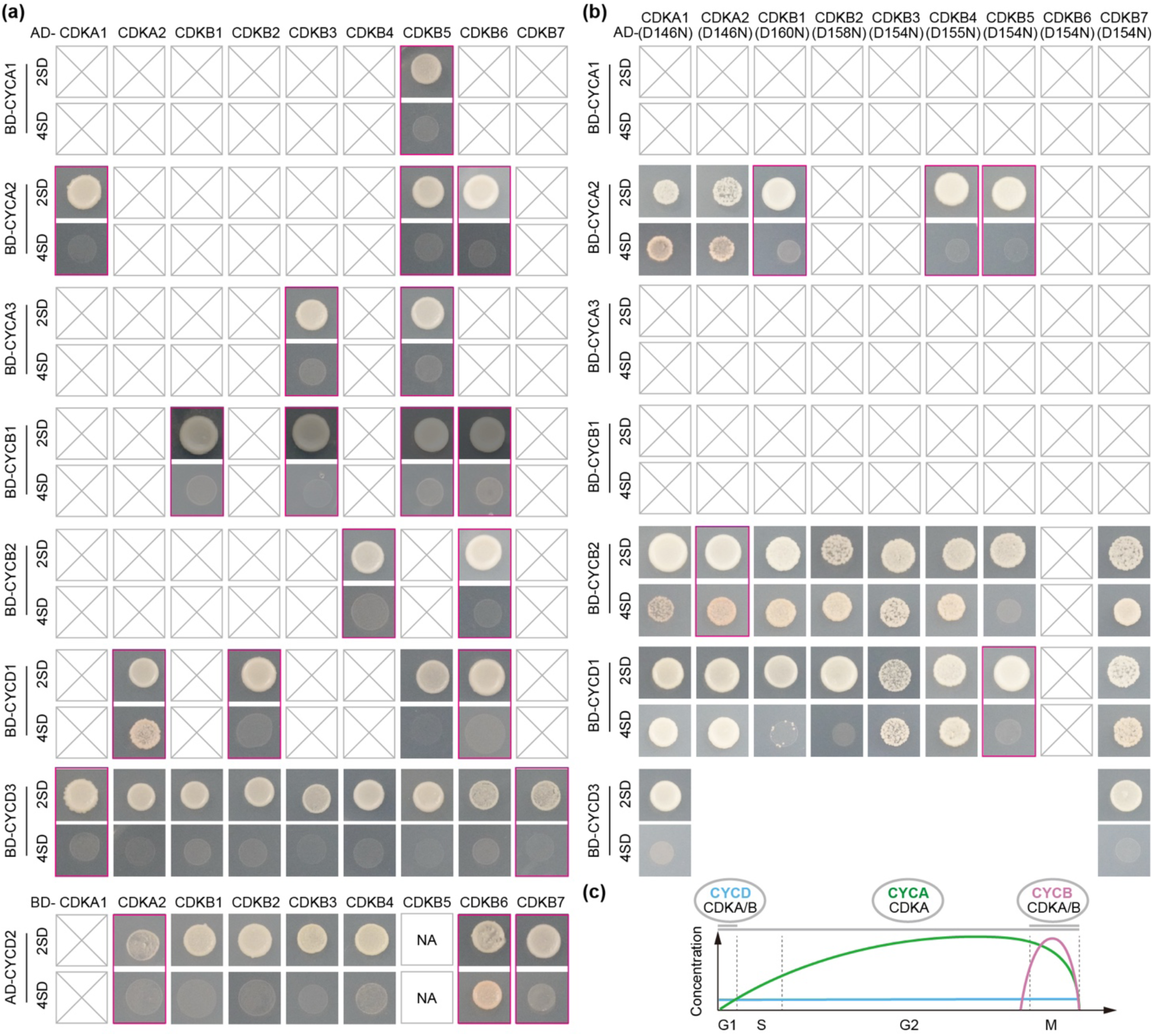
Yeast two-hybrid analysis of cyclin-CDKA/B interactions and a proposed model of cyclin-CDK function in the moss *P. patens*. (a, b) Interactions between cyclins and *WT* (a) or activity-defective (b) CDKA/B members assessed by yeast two-hybrid assays. Gray squares with diagonals indicate transformants with no viable colonies; red boxes indicate transformants with fewer than three viable colonies. NA, not available. (c) Proposed model of cyclin-CDK function in *P. patens*. Moss gametophyte cells exhibit a short G1 phase and a prolonged S/G2 phase. Cyclin D is constitutively expressed throughout the cell cycle and acts with CDKA/B members to regulate tissue growth in G1. Cyclin A is degraded during M phase and pairs with CDKA to contribute to progression through multiple cell-cycle phases. Cyclin B is specifically expressed during M phase and governs mitotic progression in complex with CDKA/B.

The deleterious effects observed in cyclin-CDK co-transformants could arise either from elevated CDK activity driven by cyclin-CDK interactions or from non-specific additive toxicity. To distinguish between these possibilities, we co-transformed cyclins with activity-defective variants of CDKA/B members. As expected, approximately 80% of these transformants (23/29), excluding CYCA1, CYCA3, and CYCB1, were viable. Among these, CYCB2 interacted with all CDKAs and CDKBs except CDKB5; CYCA2 preferentially bound CDKA1 and CDKA2; and CYCD1 associated with CDKA1, CDKA2, CDKB3, CDKB4, and CDKB7 (Figure 7b). Interestingly, co-transformation of CYCA2 with CDKB2, CDKB3, or CDKB7 remained lethal even with activity-defective variants, implying non-specific additive effects in these combinations. In addition, while wild-type CDKB6 did not affect yeast viability, its activity-defective form led to few or no colonies when transformed alone or with cyclins (Figures 7a and 7b), suggesting that activity-defective CDKB6 is toxic in yeast. Nevertheless, our results indicate that CYCB2 broadly interacts with both CDKA and CDKB, CYCA2 preferentially associates with CDKAs, and CYCD1 binds CDKAs along with a subset of CDKBs.

## Discussion

In summary, this study reveals functional divergence of the principal cyclin classes—D, A, and B in the moss *P. patens*. We provide evidence that cyclin D, cyclin A, and cyclin B play roles in G1, S-to-G2/M, and M phase regulation, respectively, with cyclin A potentially exerting overlapping functions with both cyclin D and cyclin B (Figure 7c). This divergence aligns with findings in flowering plants and underscores the evolutionary conservation of these cyclins in non-vascular land plants.

Notably, moss gametophyte cells are predominantly arrested in G2 phase (Schween et al. 2008). Our results corroborate this feature and demonstrate that it arises from a markedly abbreviated G1 phase, as evidenced by flow cytometry and post-mitotic PCNA1 expression (Figures 6b and 6c). Although G1 phase typically provides a window for macromolecule synthesis and volume expansion before division, protonema cells of *P. patens* grow continuously throughout the cell cycle—even during M phase (Figure S9). Under such conditions, mitotic preparation may not require a prolonged G1 phase. As *P. patens* exhibits exceptional regenerative capacity (Ishikawa and Hasebe 2022), the abbreviated G1 phase is likely an evolutionarily adapted feature that supports its rapid protonemal growth.

Based on the expression profiles and overexpression phenotypes, a recent study in *M. polymorpha* proposed distinct phase-specific roles for D-, A-, and B-type cyclins at G1, S, and G2/M, respectively (Romani et al. 2026). However, our results suggest partially overlapping roles among these cyclins in *P. patens*. First, cyclin D may contribute to M-phase regulation, as *cyclin D* mutants exhibited delayed division in protonemata and an increased 4C population in gametophore cells (Figures 5c-5f and 6c). The constitutive expression of *CYCD2* throughout the cell cycle and its interaction with CDKB members also support a potential contribution to M phase (Figures 2d, 2f, 2h and 7b). Second, cyclin A appears to contribute to multiple cell-cycle phases. The early cell-cycle expression of *CYCA2* and the phenotypic similarity of *cyca* mutants and overexpressors to *cycd* lines implicate CYCAs in the G1/S transition (Figures 2e-2h and 3). Moreover, CYCA2 preferentially associates with CDKA1 and CDKA2, which have previously been shown to govern G1/S transition (Bao et al. 2022) (Figure 7b). The involvement of CYCAs in G2/M is supported by findings that *CYCA2* expression declines during mitosis (Figures 2e and 2f) and the *cyca1,a2,a3* mutant exhibits a markedly elevated 4C population (Figure 6c). Lastly, CYCAs play an important role in S-to-G2/M transition, as *CYCA2* expression closely mirrors that of *PCNA1* and its ectopic expression provokes mitotic entry (Figures 6b and 6d).

Investigating cyclins and CDKs in moss enables the comparison of cell-cycle adaptation between simple (protonemata) and complex (gametophores) tissues. Protonemal tissue consists of a two-dimensional filamentous network established by iterative tip cell growth and branching, whereas cells in gametophores undergo diffuse growth and are organized in a three-dimensional body (Moody 2022, Naramoto et al. 2022). Our data suggest that these tissues exhibit differential sensitivity to cyclin perturbations. For instance, although mitotic progression is severely impaired upon *CYCB1* overexpression (Figures 5a and 5b), a substantial proportion of protonema cells were able to survive, and the overall protonema development remained largely unaffected (Figure 3). In contrast, gametophores almost completely failed to mature due to stochastic cell death at various developmental stages (Figures 4b, 4c, and 4i). This pattern suggests that loss of small cell populations—perhaps stem cells—within gametophores severely compromises subsequent organogenesis, whereas protonemata could either tolerate failed divisions or regenerate apical stem cells upon cell death, rendering them less susceptible to *CYCB1* overexpression. These findings imply that complex tissues are more sensitive to cell-cycle perturbations and depend on precise cell-cycle regulation for normal development. Future comparative studies between protonemata and gametophores could help clarify how cell-cycle regulation has been adapted to support the transition from two-dimensional to three-dimensional development.

## Materials and Methods

### Moss cultivation

Moss colonies were routinely maintained on standard BCDAT plates. To obtain large-scale protonemal tissue for transgenesis, plants were homogenized using a tissue homogenizer (Omni Inc., homogenizer cat. TH220-PCRH, tip cat. 30750H) and propagated on BCDAT plates overlaid with cellophane. For confocal imaging, a small piece of protonemal tissue was inoculated onto a thin layer of BCD medium in 35-mm glass-bottom dishes (MatTek, cat. P35G-1.5-14-C) and cultured for 5–7 days. All cultures were grown at 24°C under continuous white light. Details for moss lines used in this study are provided in Table S2.

### Sequence analyses

Cyclin and CDK sequences were retrieved from the Phytozome and UniProt databases (Goodstein et al. 2012, UniProt 2023). Transcript expression data were obtained from the eFP browser (Ortiz-Ramirez et al. 2016). Protein sequences were aligned using Clustal Omega as implemented in Jalview (Waterhouse et al. 2009, Sievers et al. 2011). Phylogenetic trees were constructed with the FASTTREE program on the Galaxy platform (Price et al. 2010, Galaxy 2026). Nuclear localization signals and PEST motifs were predicted using cNLS Mapper and EMBOSS epestfind, respectively (Rice et al. 2000, Kosugi et al. 2009). Sequences used for analysis are provided in Table S1.

### Molecular biology

The 2 × SV40 sequence (ATGGCTGATCCAAAAAAGAAGAGAAAGGTAGATCCTAAGAAGAAACGTAAGGTGGA CGGT) was commercially synthesized and inserted into the pENTR-Citrine vector immediately upstream of the Citrine coding sequence. The myristoylation (Myri) signal (ATGGGATGCTTCTCGAGTAAGCCCAAG) for membrane targeting was derived from *PpBSK-b4/Pp6c18_8140/Pp3c18_18060* (Lai et al. 2023). The F2A self-cleaving peptide coding sequence (CATAAGCAGAAAATCGTTGCACCAGTTAAGCAGACGTTGAACTTTGATTTGCTTAAA CTAGCAGGTGATGTTGAATCCAATCCTGGACCT) was obtained from foot-and-mouth disease virus 2A (Khosla et al. 2020). Both elements were inserted into the pENTR-2xSV40-Citrine plasmid via PCR-based mutagenesis or InFusion (Takara, cat. 638948) together with mCherry-FLAG coding sequence. The resulting pENTR-Myri-mCherry-F2A-2xSV40-Citrine cassette was subsequently transferred into pMY337 (a kind gift from Moé Yamada, Nagoya University) between the *EF1α* promoter and *rbcS* terminator by LR reaction (Thermo, cat. 11791019). To generate cyclin overexpression constructs, the genomic fragments of *CYCD2*, *CYCA2*, and *CYCB1* — encompassing their native promoters (1421 bp, 1682 bp, and 1246 bp, respectively) and complete coding regions—were inserted into the pENTR vector upstream of the mCherry-FLAG-F2A-2×SV40-Citrine sequence, replacing the myristoylation signal via InFusion. These expression cassettes were subsequently transferred into the pPY98 Gateway vector upstream of the *nos* terminator by LR reactions. The *CYCA2(NT)* overexpression construct was generated from the *CYCA2-ox* plasmid by deleting the region encoding CYCA2(204-480) through mutagenesis. To create the *VPS4B-CYCB1(NT)-mNeonGreen* overexpression construct, the genomic region encoding CYCB1(1-251) was inserted into pENTR-VPS4B-mNG using InFusion, and the entire cassette was subsequently transferred into pMY337. For CRISPR-based editing, single-guide RNA (sgRNA) targets were manually designed and cloned into pPY156 via mutagenesis; target sequences and oligonucleotide templates are provided in Figure S6. For inducible expression, the genomic coding region of *CYCD2* and the coding sequence (CDS) of *CYCA2* were fused in-frame with a C-terminal Cerulean (CR) or mNG tag, respectively, within the pENTR vector. These cassettes were subsequently transferred into pPGX8 via LR reactions (Kubo et al. 2013). For endogenous *PCNA1* labeling, the mNG-FLAG-F2A-2×SV40-mCherry coding sequence was inserted between two homology arms via InFusion: one arm consisted of a 1061-bp fragment immediately upstream of the *PCNA1* stop codon, and the other consisted of a 1622-bp fragment downstream of the stop codon. Yeast two-hybrid plasmids were constructed by inserting the CDS of each cyclin or CDK gene into pGADT7 and pGBKT7 vectors via InFusion. Activity-defective variants were generated by introducing the corresponding mutations into the wild-type yeast plasmids through mutagenesis. All primers and plasmids are provided in Table S3 and Table S4.

### Moss transformation

Protonemata harvested one week after subculture were digested with 0.5% (w/v) driselase. The released protoplasts were washed with 8% mannitol and resuspended in MMM buffer (9% mannitol, 0.1% MES pH 5.6, 15 mM MgCl_2_). For overexpression and fluorescent protein knock-in lines, 300 μL of protoplasts were mixed with 30 μg of linearized targeting plasmid and incubated with 300 μL of PEG6000 solution at room temperature for 30 min. The mixture was then supplemented with 1.5 ml of pre-chilled W5 buffer (2 mM MES pH 5.7, 5 mM KCl, 125 mM CaCl_2_, 154 mM NaCl) and centrifuged to collect the protoplasts. Protoplasts were resuspended in PRMT medium, plated onto cellophane-overlaid PRM plates, and cultured under standard conditions. For *PCNA1* endogenous labeling, the knock-in construct was co-transformed with a Hygromycin B-resistance plasmid. For CRISPR/Cas9 editing, 10 μg of sgRNA plasmid was co-transformed with 10 μg of pAct-Cas9 plasmid; to generate *CYCB1* hypomorphic mutants, 10 μL of 50 μM oligonucleotide templates were additionally included. Transformants were selected on appropriate medium, once for genome editing and *PCNA1* knock-in lines, and twice for overexpression lines. Positive colonies were subcultured and validated by PCR and DNA sequencing.

### Staining and chemical treatment

To visualize protonema cell morphology, protonemata cultured in 35-mm dishes were stained with 10 µg/ml calcofluor white (Sigma, cat. 18909-100ML-F) for 10 min and imaged within 1 h. For morphological analysis of gametophore buds or young gametophores, samples were stained with 30 μM propidium iodide (Sigma, cat. P4864) for 10 min prior to imaging. For inducible expression of *CYCD2* and *CYCA2*, protonemata were treated with 1 µM β-estradiol (Coolaber, cat. CE5114) for 10 min and imaged 1 h post-treatment. Gametophore development was induced by a 10-min treatment with 1 μM 2iP (Nacalai Tesque, cat. 02744-94); tissues were then imaged at the defined developmental stages, specifically early buds (24–48 h) and late buds/young gametophores (48–96 h). For cytokinin-induced transcript analysis, protonemata were harvested 36 h post-2iP application and total RNA was extracted (Oriscience, cat. NB304) for real-time quantitative PCR analysis.

### Confocal microscopy and image processing

High-resolution imaging data were acquired using an Olympus SpinSR spinning-disk confocal system, with operations managed via cellSens Dimension software (version 4.3.1). The key hardware elements consisted of an IX83P2ZF inverted microscope stand featuring a series of objectives (10×/NA 0.40, 20×/NA 0.70, 40×/NA 0.95, 60×/NA 1.42, and 100×/NA 1.48), a Yokogawa CSU-W1 confocal scanner unit utilizing 405-, 488-, and 561-nm diode lasers, and a Hamamatsu ORCA Flash 4.0 V3 sCMOS camera for signal acquisition. Laser output was fixed at 50% for all wavelengths. Exposure parameters were 500 ms for the 488- and 561-nm channels and 100 ms for the 405-nm channel. Time-lapse series were recorded at 2-min intervals. Images were processed and analyzed using Fiji software (version 2.14.0). Because tip-cell length varies during the cell cycle, we quantified the length and width of the first subapical cell. To analyze cell-cycle phase durations, we measured three distinct time intervals: (i) from nuclear envelope breakdown (NEB) to metaphase completion (NEB-to-meta), (ii) from anaphase onset to nuclear envelope reformation (anaphase), and (iii) from the initiation of phragmoplast expansion to its eventual disappearance (cytokinesis).

### Flow cytometry

Freshly cultured protonemata, gametophores, or *Arabidopsis* flowers were transferred to 1.5-mL tubes, flash-frozen in liquid nitrogen, and mechanically disrupted into a coarse powder. The frozen material was resuspended in 600 μL of ice-cold TMT buffer (200 mM Tris-HCl pH8.0; 4 mM MgCl_2_; 0.5% Triton X-100; 16.7 μg/mL RNase A). The resulting cell lysates were passed through a 35-µm cell strainer (Falcon, cat. 352235) and then incubated at 37°C for 10 min. Each lysate was supplemented with 30 µL of 1 mg/mL propidium iodide (Sigma, cat. P4170), and then stained at 37°C for an additional 10 min. The stained suspensions were filtered once more through a 35-µm cell strainer cap and collected in 5 mL round-bottom polystyrene tubes (Falcon, cat. 352235). Samples were analyzed on a BD LSRFortessa cell analyzer (BD Biosciences) using the following voltage settings: FSC = 730, SSC = 300, and PE-Texas Red = 345, with a PE threshold of 200. Unstained samples were processed in parallel and served as negative controls for gating.

### Yeast two-hybrid assay

Competent AH109 yeast cells were co-transformed with approximately 0.1 µg of each AD- or BD-fusion construct, supplemented with 100 µg of carrier DNA. Primary transformants were selected on agar plates deficient in tryptophan and leucine (2SD). Colonies that grew were then picked, propagated, and spotted onto medium lacking tryptophan, leucine, histidine, and adenine (4SD) to evaluate pairwise interactions. For each transformation, a minimum of three independent colonies were included in the assays, except when fewer than three viable colonies were available; any deviations from this standard are indicated in the figure legends.

### Statistical analysis

Graphical data show individual points and are presented as mean ± SD or as box plots displaying the median (or mean) and range (minimum to maximum). Sample sizes are indicated in the figure or main text. Statistical significance was assessed using the tests described in the corresponding figure legends. *P* < 0.05 was considered statistically significant.

### Accession numbers

*CYCA1 (Pp6c17_12130 / Pp3c17_22710), CYCA2 (Pp6c2_4910 / Pp3c2_8770), CYCA3 (Pp6c1_20520 / Pp3c1_39100), CYCB1 (Pp6c9_9380 / Pp3c9_18910), CYCB2 (Pp6c15_10680 / Pp3c15_21520), CYCD1 (Pp6c15_8720 / Pp3c15_17470), CYCD2 (Pp6c9_4230 / Pp3c9_8300), CYCD3 (Pp6c22_8510 / Pp3c22_17100), CDKA1 (Pp6c3_7680 / Pp3c3_15290), CDKA2 (Pp6c2_10510 / Pp3c2_18480), CDKB1 (Pp6c16_9080 / Pp3c16_3910), CDKB2 (Pp6c25_1510 / Pp3c27_6070), CDKB3 (Pp6c14_7150 / Pp3c14_12680), CDKB4 (Pp6c1_16330 / Pp3c1_31370), CDKB5 (Pp6c1_2790 / Pp3c1_5080), CDKB6 (Pp6c11_5940 / Pp3c11_10380), CDKB7 (Pp6c7_10000 / Pp3c7_18430), PCNA1 (Pp6c12_8960 / Pp3c12_18160)*.

## Data availability

All data supporting the findings of this study are available within the manuscript and its supplementary materials.

## Acknowledgements

Confocal imaging and flow cytometry were supported by the Life Science Core Facilities at the College of Life Sciences, Sichuan University. We thank Linzhu Li for imaging support, and Xiaoning Wang (Flow Cytometry Laboratory Unit, National University of Singapore) and Xiaoyang Li for technical support with flow cytometry. Financial support was provided by the Organization Department of the CPC Central Committee (Grant No. SQ2024QB02593) and the National Natural Science Foundation of China (Grant No. 32100273) to P.Y.

## Author Contributions

Conceptualization: P.Y.; Molecular biology: Y.G., M.L.; Transgenesis: Y.G., J.R.; Imaging: Y.G., J.R., P.Y.; Y2H: Y.G., M.L.; Flow cytometry: Y.G., C.L., P.Y.; Data analysis: Y.G., P.Y.; Writing and proofreading: Y.G., J.R., P.Y.

## Competing interests

The authors declare no competing interests.

**Table S1. Complete list of cyclin and CDK sequences analyzed in this study.**

**Table S2. Moss strains used in this study.**

**Table S3. Primers used in this study.**

**Table S4. Plasmids used in this study.**

**Movie S1. CYCD2 expression in a protonema cell.**

**Movie S2. CYCA2 expression in a protonema cell.**

**Movie S3. CYCA2 N-terminal domain expression in a protonema cell.**

**Movie S4. CYCB1 expression in a protonema cell.**

**Movie S5. VPS4A localization in a protonema cell.**

**Movie S6. VPS4B-CYCB1(NT) fusion protein localization in a protonema cell.**

**Movie S7. Normal cell division in the *CYCB1* overexpressor.**

**Movie S8. Defective cell division in the *CYCB1* overexpressor.**

**Movie S9. Mitotic division of a wild-type protonema cell.**

**Movie S10. Mitotic division of a *cycd1,d2,d3* triple mutant cell.**

**Movie S11. Mitotic division of a *cyca1,a2,a3* triple mutant cell.**

**Movie S12. Mitotic division of a *cycb1^L357H^,b2* double mutant cell.**

**Movie S13. PCNA1 expression and localization in a protonema cell.**

---

**Figure S1.**
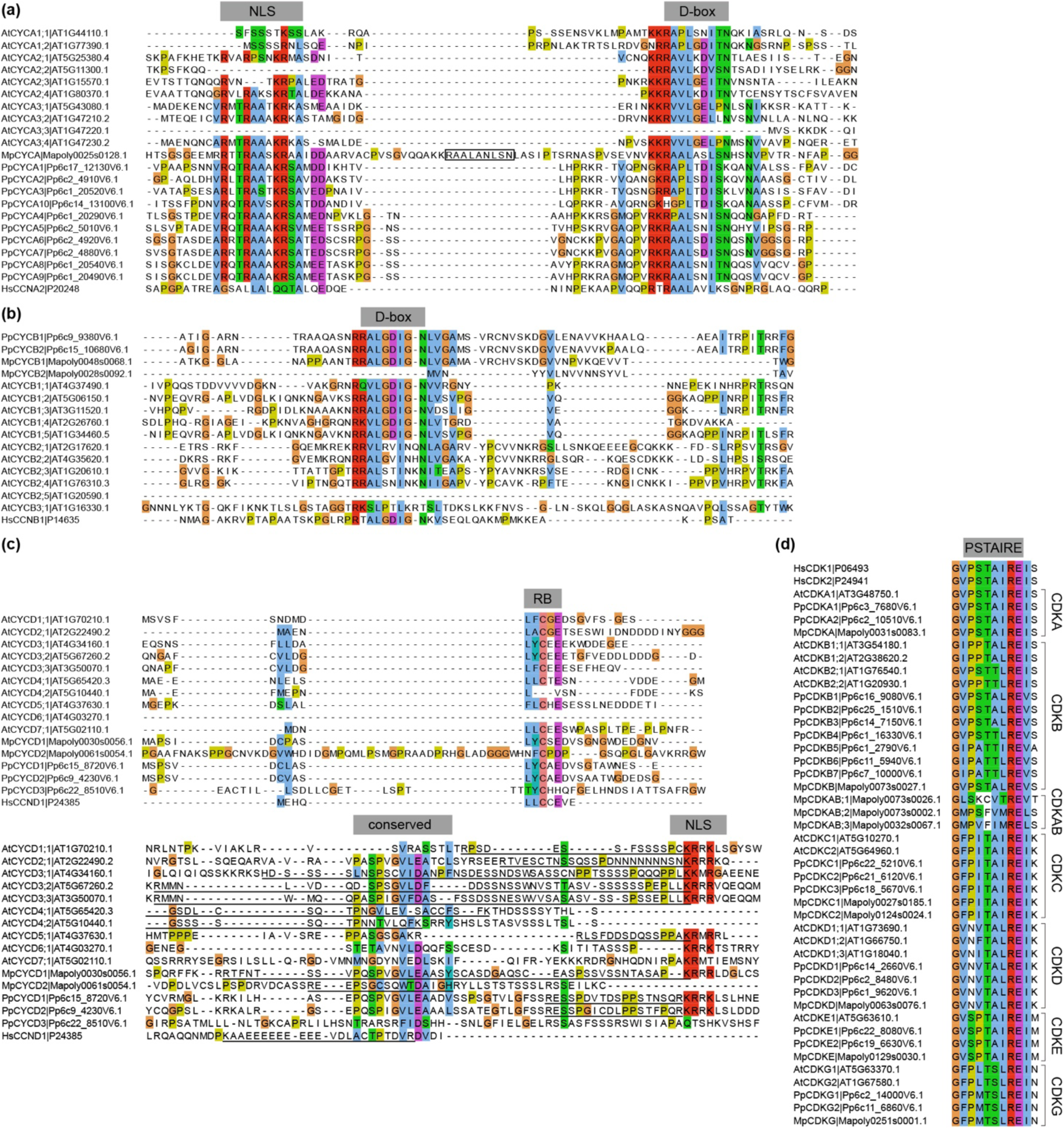
Sequence alignment of cyclins and CDKs. (a-c) Alignment of conserved regulatory regions in A-(a), B-(b), and D-type (c) cyclins. An additional D-box (boxed region) is present in the liverwort CYCA sequence. Predicted PEST motifs at the C-termini of CYCDs are underlined (shown as partial sequences for AtCYCD4;1 and AtCYCD4;2). D-box, destruction box; NLS, nuclear localization signal; RB, Retinoblastoma-related protein-binding motif. (d) Alignment of PSTAIRE motifs in CDKs.

**Figure S2.**
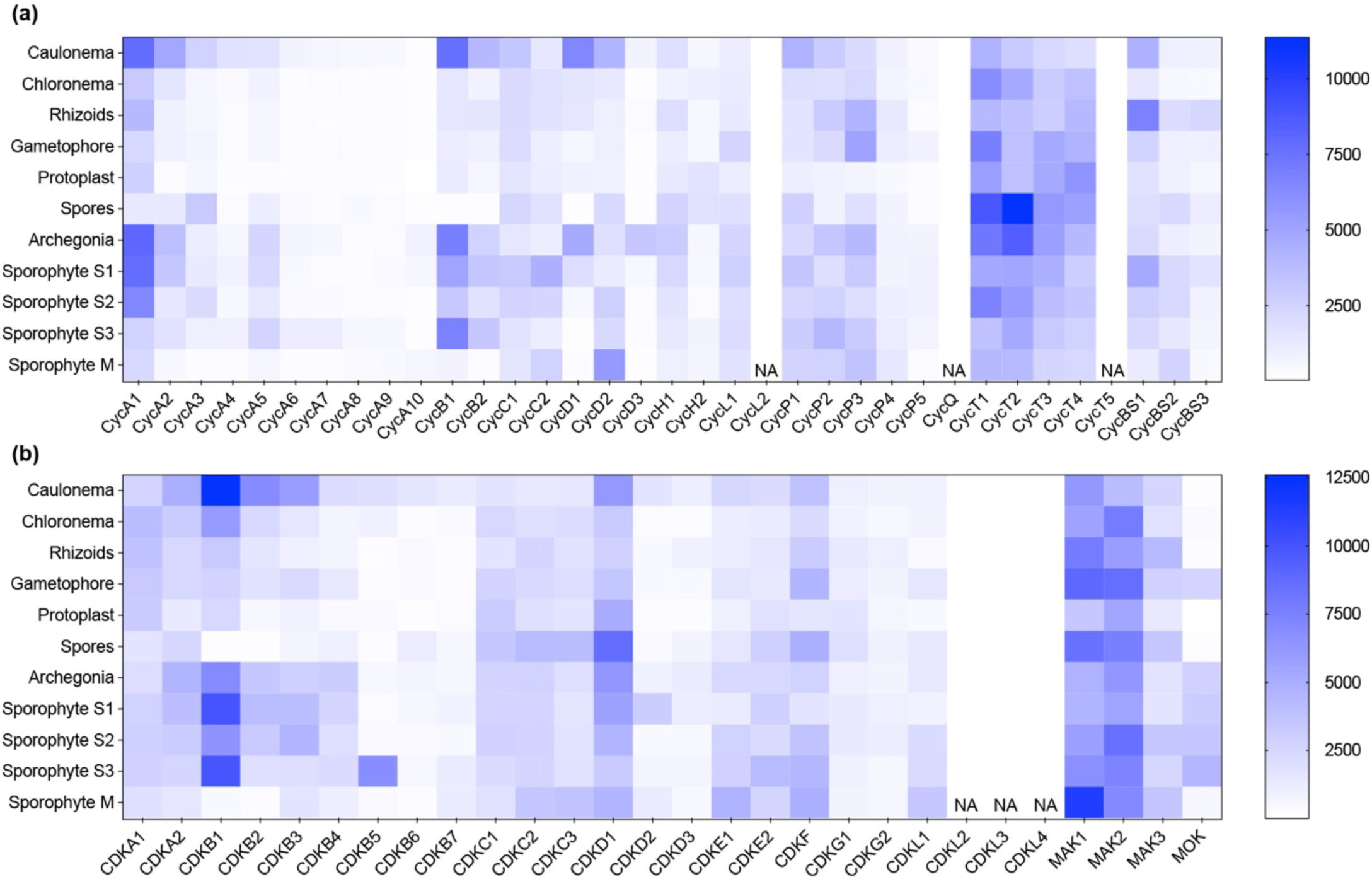
Transcript levels of moss cyclins and CDKs. (a, b) Expression levels of cyclins (a) and CDKs (b) across major tissues and cell types. NA, not available.

**Figure S3.**
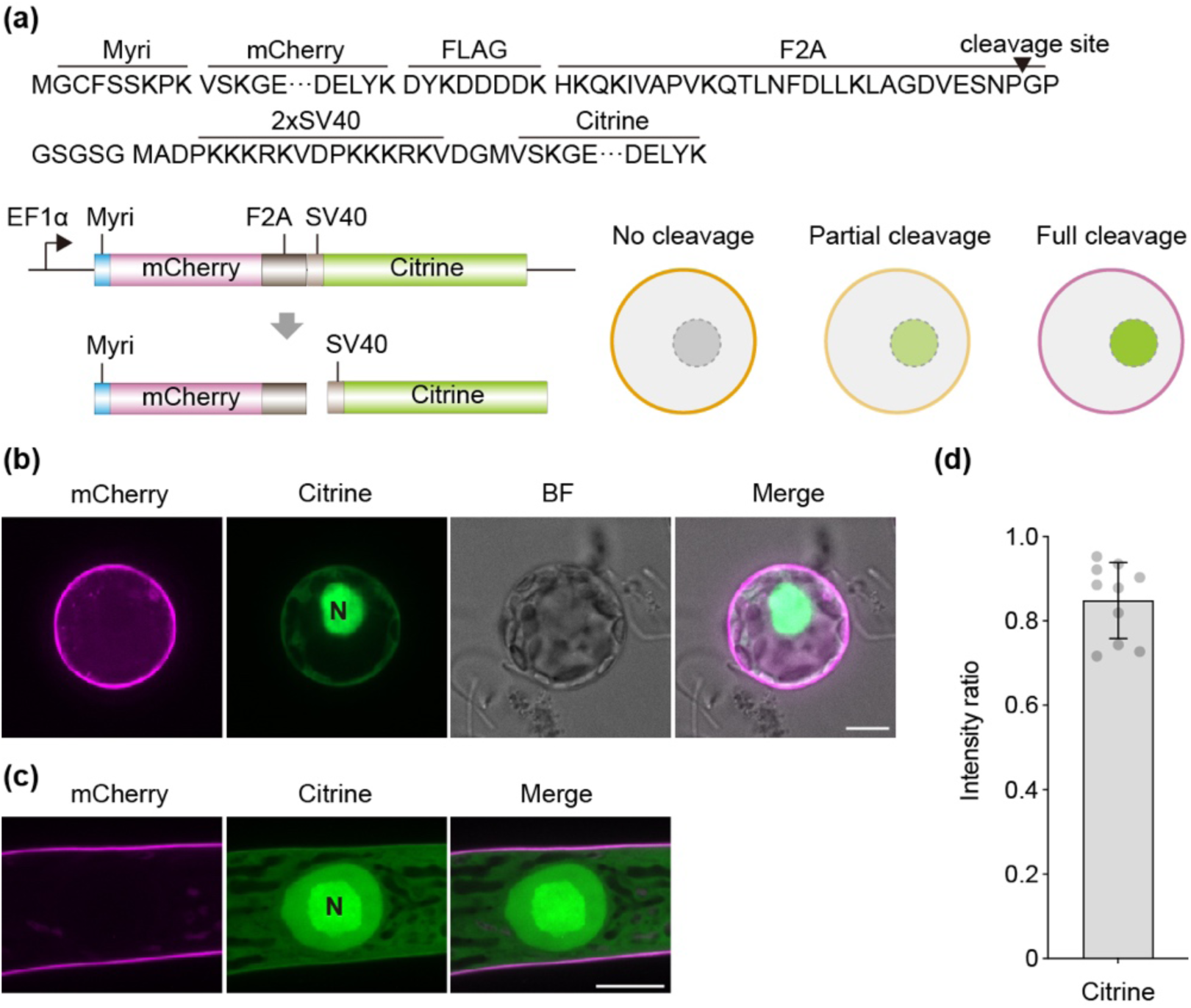
Analysis of F2A cleavage efficiency in moss. (a) Schematic of the F2A self-cleaving reporter construct, consisting of an N-terminal myristoylation signal (Myri) fused to mCherry-FLAG and a 2×SV40-Citrine fragment downstream of the F2A peptide. The myristoylation signal directs membrane targeting, whereas SV40 enables nuclear localization. (b, c) Expression of the F2A reporter in protoplasts (b) and stably transformed protonema cells (c). BF, bright field; N, nucleus. Scale bars, 10 μm. (d) F2A cleavage efficiency calculated as the nuclear-to-total intensity ratio (nuclear / [membrane + nuclear]). Since SV40-Citrine is partially cytoplasmic and its membrane-associated signal cannot be unambiguously distinguished from cytoplasmic fluorescence, the measured efficiency is likely underestimated. Data are shown as mean ± SD (n = 10).

**Figure S4.**
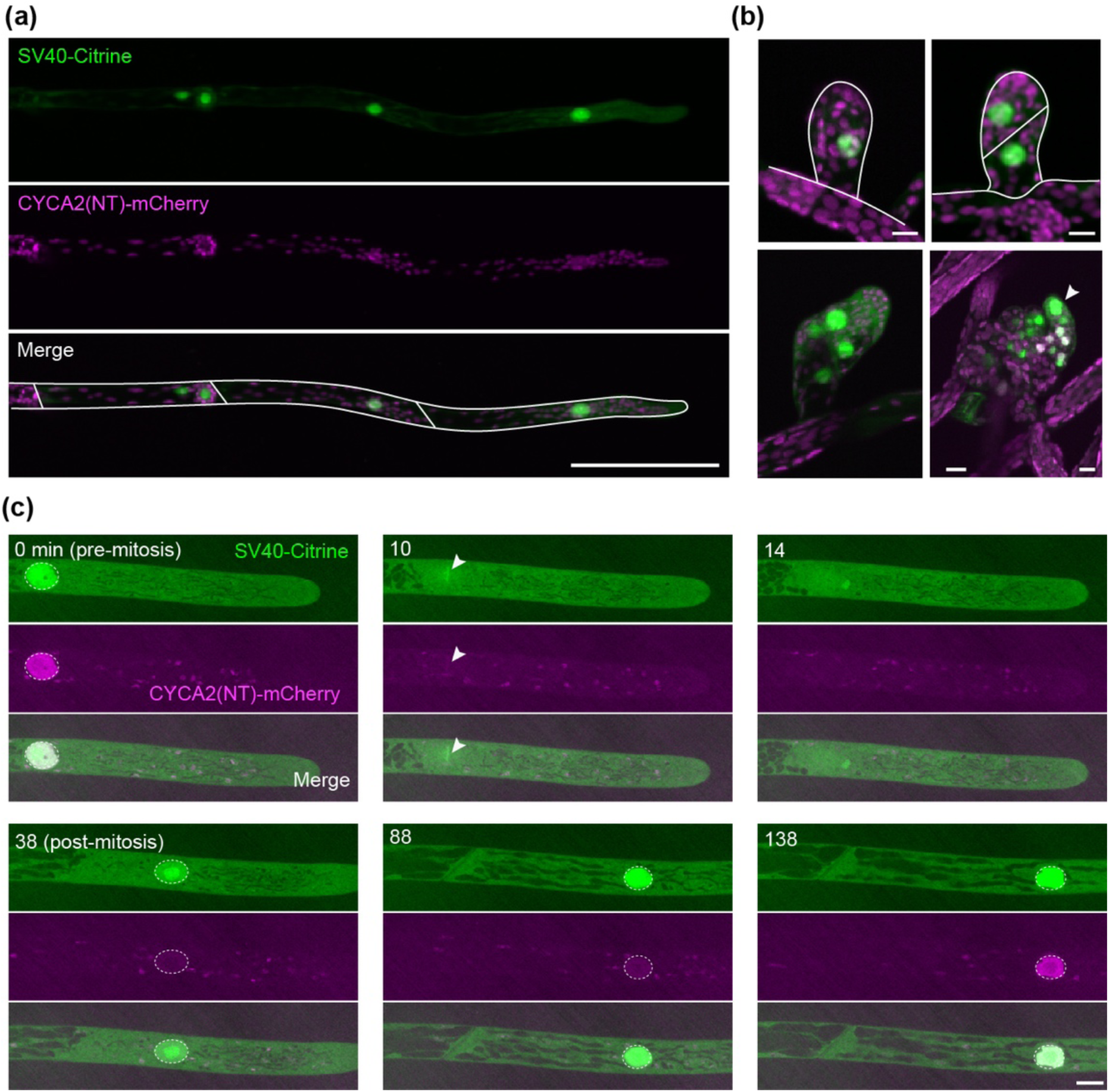
Expression and localization of the CYCA2 N-terminal domain. (a) Expression of CYCA2(NT)-mCherry-F2A-SV40-Citrine in protonema cells. Scale bar, 100 μm. (b) Expression of CYCA2(NT) in young gametophores. The arrowhead indicates a phyllid initial cell. Scale bars, 10 μm. (c) Time-lapse imaging of CYCA2(NT) during protonema cell division. Arrowheads indicate chromosomes; nuclei are outlined by dashed circles. Scale bar, 10 μm.

**Figure S5.**
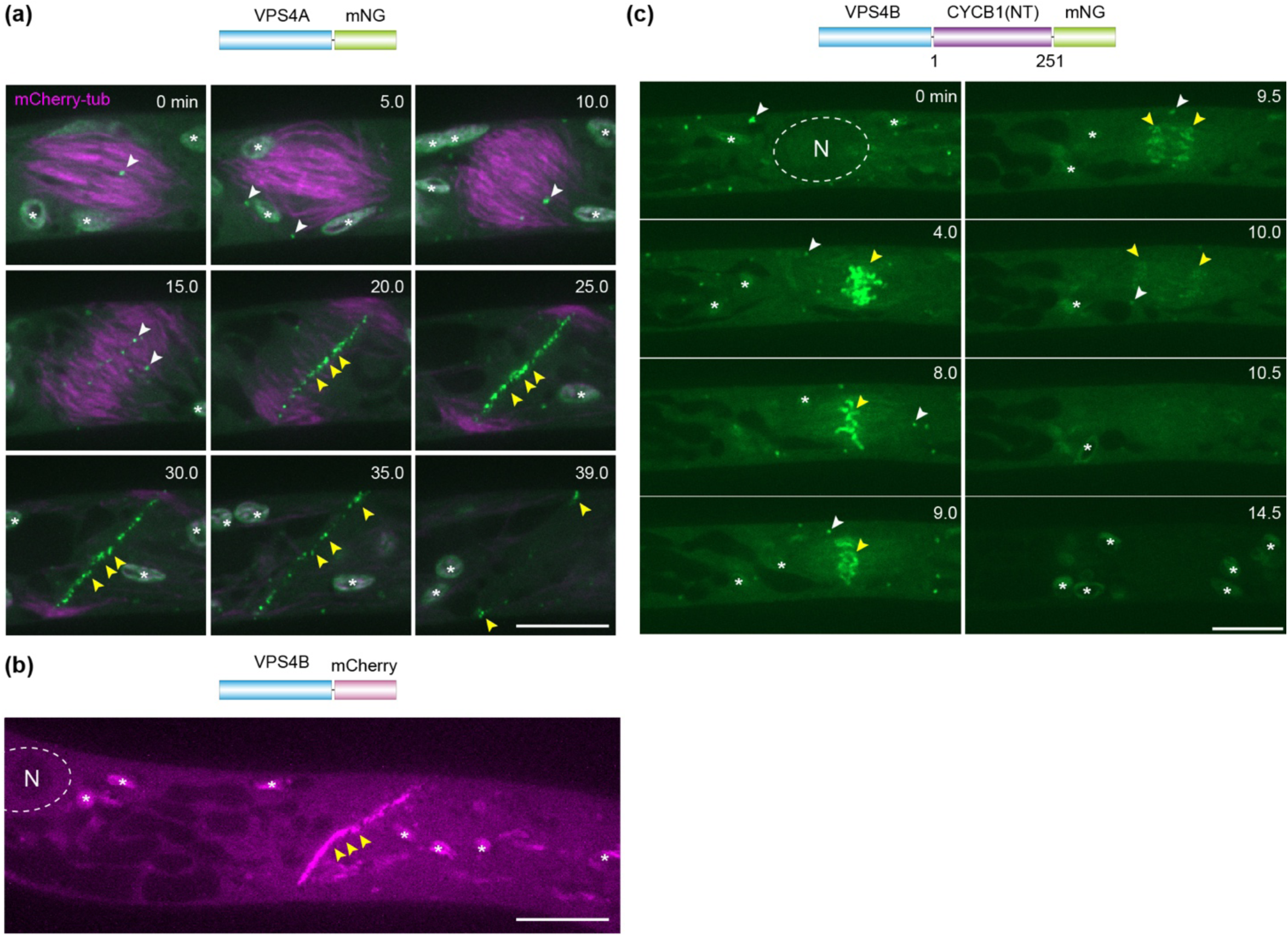
The CYCB1 N-terminal domain targets VPS4B to mitotic chromosomes and promotes its degradation. (a) During cytokinesis in wild-type cells, VPS4A-mNG decorates endosomes (white arrowheads) and the cell plate (yellow arrowhead). The mCherry-tubulin reporter marks the spindle and phragmoplast. Asterisks indicate chloroplast autofluorescence. Scale bar, 10 μm. (b) Like VPS4A, VPS4B enriches at the cell plate. The nucleus is demarcated by a dashed circle. Scale bar, 10 μm. (c) The VPS4B-CYCB1(NT) fusion protein labels endosomes (white arrowheads) but is redirected to chromosomes during mitosis (yellow arrowheads), followed by rapid degradation. Scale bar, 10 μm.

**Figure S6.**
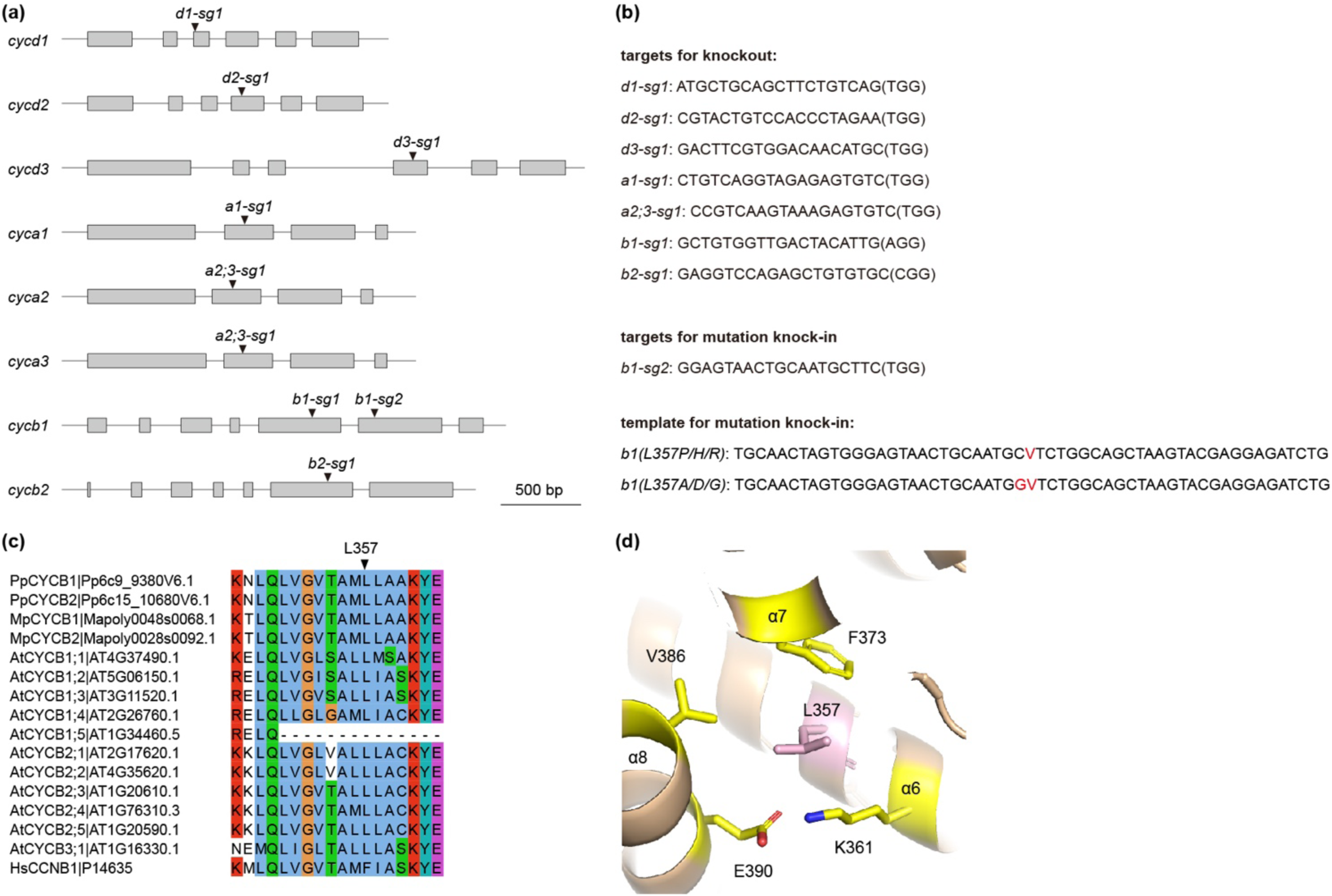
Generation of cyclin mutants using CRISPR/Cas9. (a) Schematic of cyclin gene structures showing the locations of CRISPR targets. Exons are depicted as gray boxes. (b) Sequences of CRISPR targets and oligonucleotide templates. Protospacer-adjacent motifs (PAMs) are indicated in brackets; point mutations in the templates are highlighted in red. (c) Sequence alignment of cyclin B members highlighting the conserved leucine residue at position 357. (d) Position of Leu357 within a hydrophobic core of the AlphaFold3-predicted structure of *P. patens* CYCB1.

**Figure S7.**
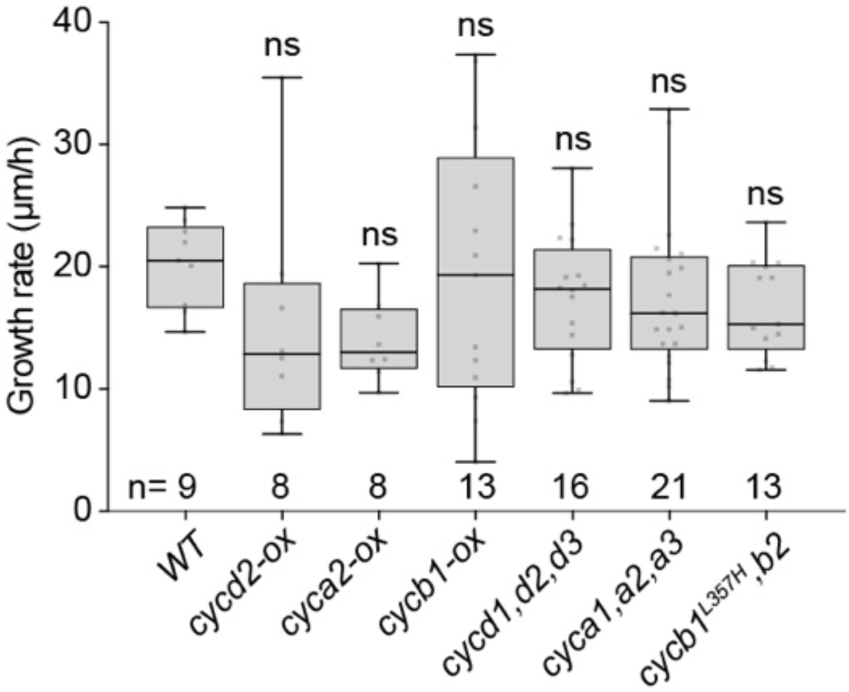
Quantification of tip cell growth rates. Sample sizes are indicated below the bars. Statistical significance relative to *WT* was assessed using Kruskal–Wallis followed by Dunn’s post hoc multiple-comparison test; ns, not significant.

**Figure S8.**
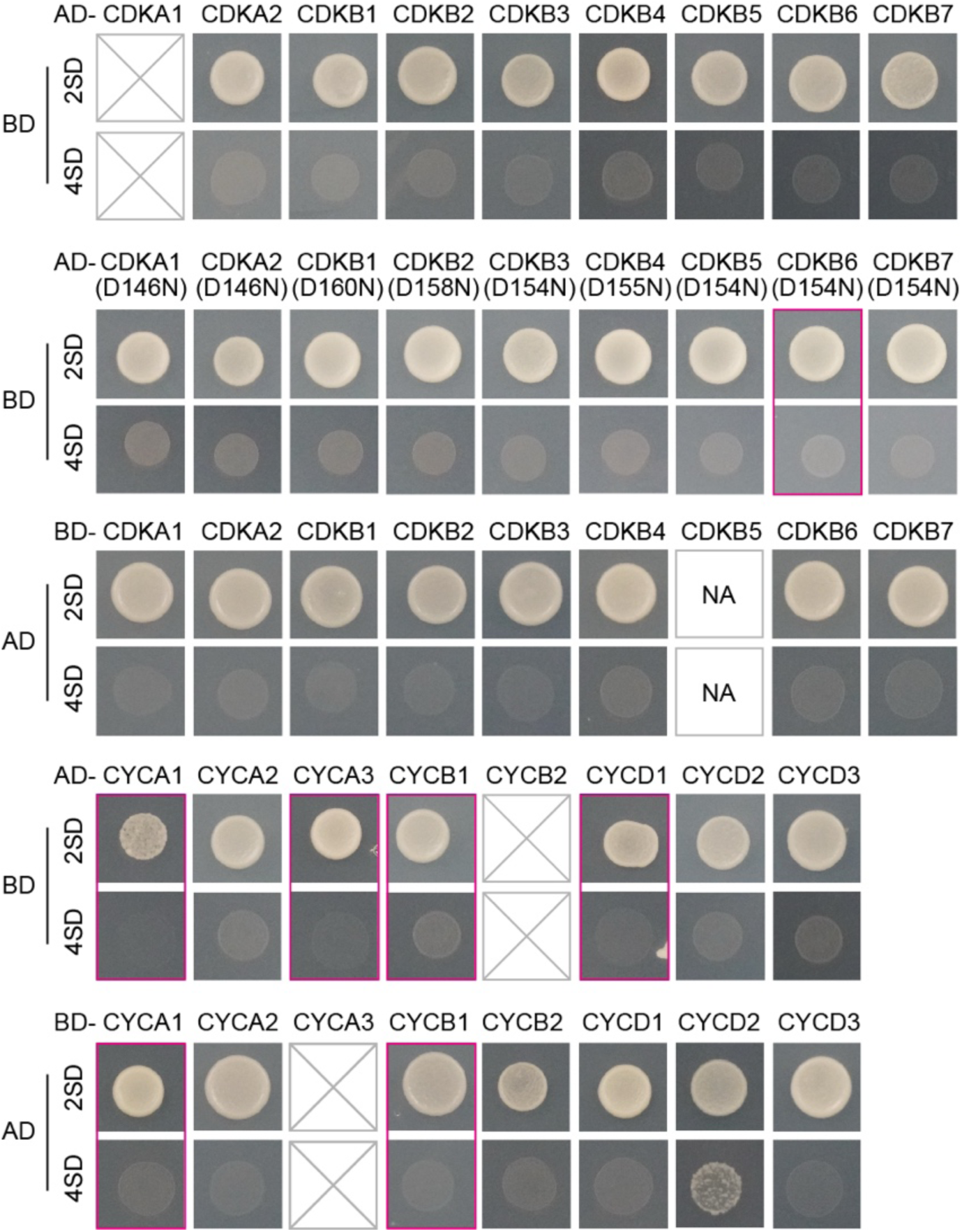
Assessment of cyclin and CDK autoactivation in yeast two-hybrid assays. Gray squares with diagonals indicate transformants with no viable colonies; red boxes indicate transformants with fewer than three viable colonies. NA, not available.

**Figure S9.**
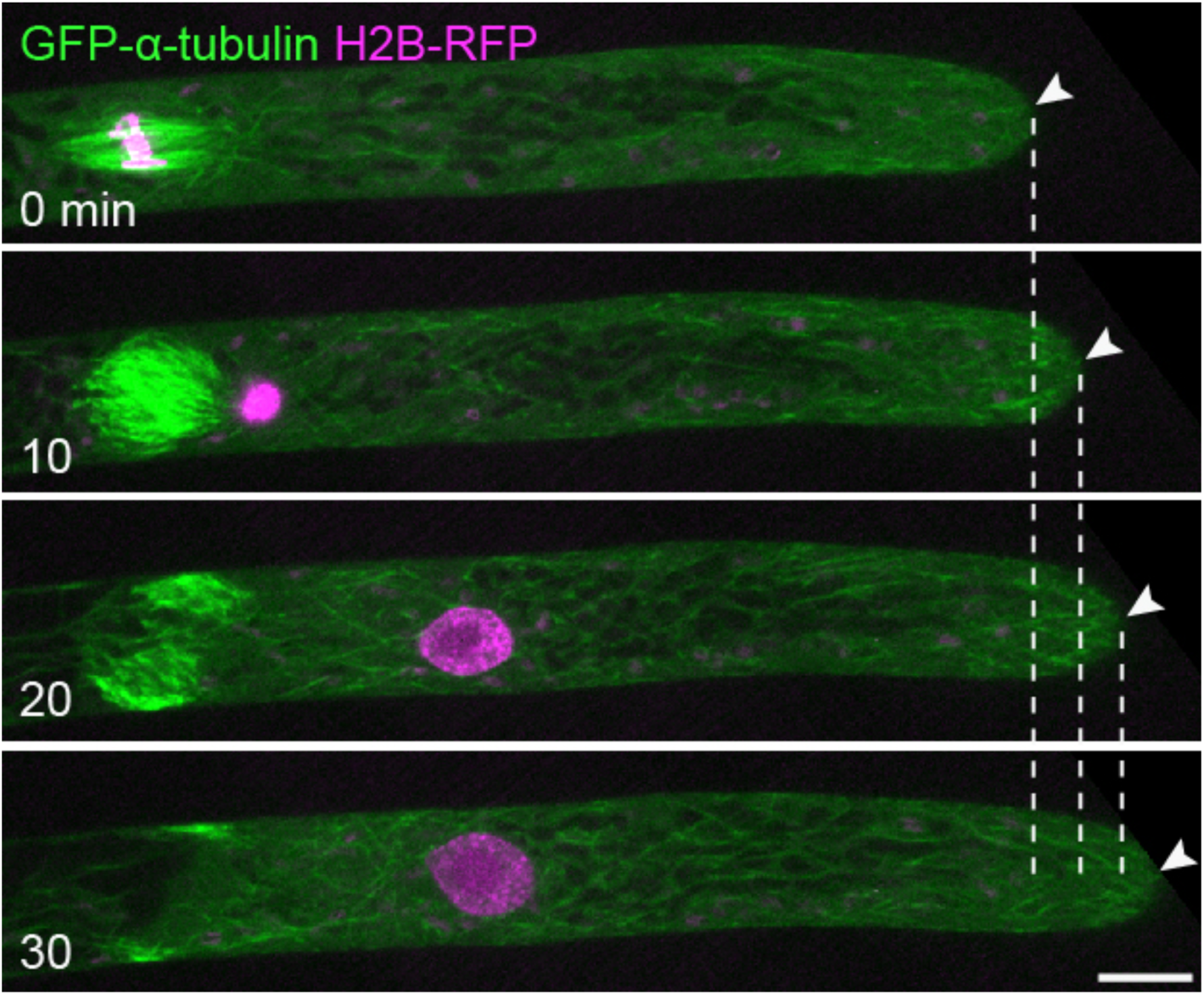
The protonema cell undergoes continuous apical growth during M phase. Time-lapse images of a dividing protonema cell. The growing apex is marked by white arrowheads. Dashed lines indicate the relative position of the apex over time. Scale bar, 10 μm.

